# Development of highly potent non-covalent inhibitors of SARS-CoV-2 3CLpro

**DOI:** 10.1101/2022.08.10.503531

**Authors:** Ningke Hou, Lei Shuai, Lijing Zhang, Xuping Xie, Kaiming Tang, Yunkai Zhu, Yin Yu, Wenyi Zhang, Qiaozhu Tan, Gongxun Zhong, Zhiyuan Wen, Chong Wang, Xijun He, Hong Huo, Haishan Gao, You Xu, Jing Xue, Chen Peng, Jing Zou, Craig Schindewolf, Vineet Menachery, Wenji Su, Youlang Yuan, Zuyuan Shen, Rong Zhang, Shuofeng Yuan, Hongtao Yu, Pei-Yong Shi, Zhigao Bu, Jing Huang, Qi Hu

## Abstract

The SARS-CoV-2 virus is the causal agent of the ongoing pandemic of coronavirus disease 2019 (COVID-19). There is an urgent need for potent, specific antiviral compounds against SARS-CoV-2. The 3C-like protease (3CLpro) is an essential enzyme for the replication of SARS-CoV-2 and other coronaviruses, and thus is a target for coronavirus drug discovery. Nearly all inhibitors of coronavirus 3CLpro reported so far are covalent inhibitors. Here, we report the development of specific, non-covalent inhibitors of 3CLpro. The most potent one, WU-04, effectively blocks SARS-CoV-2 replications in human cells with EC_50_ values in the 10-nM range. WU-04 also inhibits the 3CLpro of SARS-CoV and MERS-CoV with high potency, indicating that it is a pan-inhibitor of coronavirus 3CLpro. WU-04 showed anti-SARS-CoV-2 activity similar to that of PF-07321332 (Nirmatrelvir) in K18-hACE2 mice when the same dose was administered orally. Thus, WU-04 is a promising drug candidate for coronavirus treatment.

**One-Sentence Summary:** A oral non-covalent inhibitor of 3C-like protease effectively inhibits SARS-CoV-2 replication.

## Main Text

The ongoing pandemic of coronavirus disease 2019 (COVID-19), which is caused by severe acute respiratory syndrome coronavirus 2 (SARS-CoV-2) (1,2), is becoming a long-term threat to human health. Vaccines and antivirals are two essential countermeasures for fighting viral infectious diseases. Despite the approval and deployment of several vaccines, millions around the world are still being infected by SARS-CoV-2 every week. SARS-CoV-2 variants that render the currently approved vaccines less effective have emerged (3–6). Antiviral agents against SARS-CoV-2 are needed. Several monoclonal antibodies, which target the spike protein of SARS-CoV-2 to block viral entry into human cells, have received an Emergency Use Authorization for treating COVID-19 {Corti:2021fp}, however, viral variants with mutations in the spike protein have been found to compromise the efficacy of these antibodies (7,8). In addition, the high cost in the production and delivery of monoclonal antibodies hinders their wide use. Potent small molecule inhibitors of SARS-CoV-2 are needed and have become the focal point of SARS-CoV-2 drug development.

The 3C-like protease (3CLpro) is an established target of antivirals against coronaviruses. More than 70% of the SARS-CoV-2 RNA genome encodes two polyproteins, pp1a and pp1ab, which undergo proteolytic cleavage to generate 16 non-structural proteins (nsps) (1,9). The cleavage is catalyzed by two proteases: nsp3 and nsp5 (also known as 3CLpro or the main protease). The multidomain protein nsp3 contains a papain-like protease (PLpro) domain and cleaves the peptide bonds to release nsp1, nsp2, and nsp3 (9,10). 3CLpro cleaves peptide bonds to release nsp4 to nsp16 (9). After releasing itself from pp1a or pp1ab, 3CLpro forms a homodimer with increased protease activity. Inhibition of the protease activity of 3CLpro blocks the release of nsp4 to nsp16 that are essential for coronavirus replication.

3CLpro is a cysteine protease. Its catalytic dyad consists of His41 and Cys145. Most reported inhibitors targeting SARS-CoV-2 3CLpro are covalent compounds derived from peptidic scaffolds that have an electrophile to react with the catalytic cysteine (Cys145) (11–23). One such covalent inhibitor, PF-07321332 (Nirmatrelvir), has been approved for the treatment of COVID-19 patients (24). Covalent inhibitors offer prolonged duration of inhibition, but suffer from potential side effects due to off-target reactions (25,26). Some non-covalent inhibitors were also reported, but no or modest inhibitory activity was observed (11,27–30); recently a non-covalent oral SARS-CoV-2 3CLpro inhibitor S-217622 was reported as a drug candidate for treating COVID-19 (31). In this study, we identified a novel class of potent non-covalent inhibitors of SARS-CoV-2 3CLpro. One of them blocked SARS-CoV-2 replication in human cell lines with half-maximal effective concentrations (EC_50_s) in the 10-nM range, and showed promising antiviral activity in mouse models.

## Results

### Screening of non-covalent inhibitors of SARS-CoV-2 3CLpro

DNA encoded library (DEL) technology is a powerful tool to identify small molecule binders of target proteins (32,33). We screened DELs of more than 49 billion compounds using His_6_-tagged purified recombinant SARS-CoV-2 3CLpro bound to Ni^2+^-NTA magnetic beads. Since adding extra residues at either the N- or C-terminus of SARS-CoV-2 3CLpro decreased its enzymatic activity (34), we rationally engineered the His_6_-tag into 3CLpro without compromising the protease activity. Analysis of the crystal structures of SARS-CoV-2 3CLpro revealed that residues Gly215 to Thr225 are located in a loop far from both the catalytic site and the dimer interface. We thus inserted an internal His_6_-tag between Arg222 and Phe223 of 3CLpro. As determined by a fluorescence-based assay (35), the protease activity of this internally His_6_-tagged 3CLpro (termed 3CLpro-mHis) was similar to that of the wild-type 3CLpro (WT 3CLpro) (fig. S1).

3CLpro-mHis was immobilized on Ni^2+^-NTA magnetic beads and incubated with the DEL. The bound DNA-encoded compounds were decoded by qPCR followed by DNA sequencing. DEL hits were classified by chemotypes and ranked with docking calculations. Compounds containing an isoquinoline ring and a bromophenyl ring were identified as potential binders for SARS-CoV-2 3CLpro. The off-DNA version of five such compounds were synthesized (Fig. 1a); their ability to inhibit purified SARS-CoV-2 3CLpro was evaluated. Four of these compounds inhibited 3CLpro with IC_50_s of less than 1 µM (Fig. 1b). Compounds WU-02 and WU-04 were the most potent, with IC_50_s of 71 and 72 nM, respectively. Consistent with the IC_50_ value, the isothermal titration calorimetry (ITC) data showed a high binding affinity between WU-04 and SARS-CoV-2 3CLpro, with a dissociation constant (*K*_d_) of 37 nM and 1:1 binding stoichiometry (Fig. 1c).

**Fig. 1.**
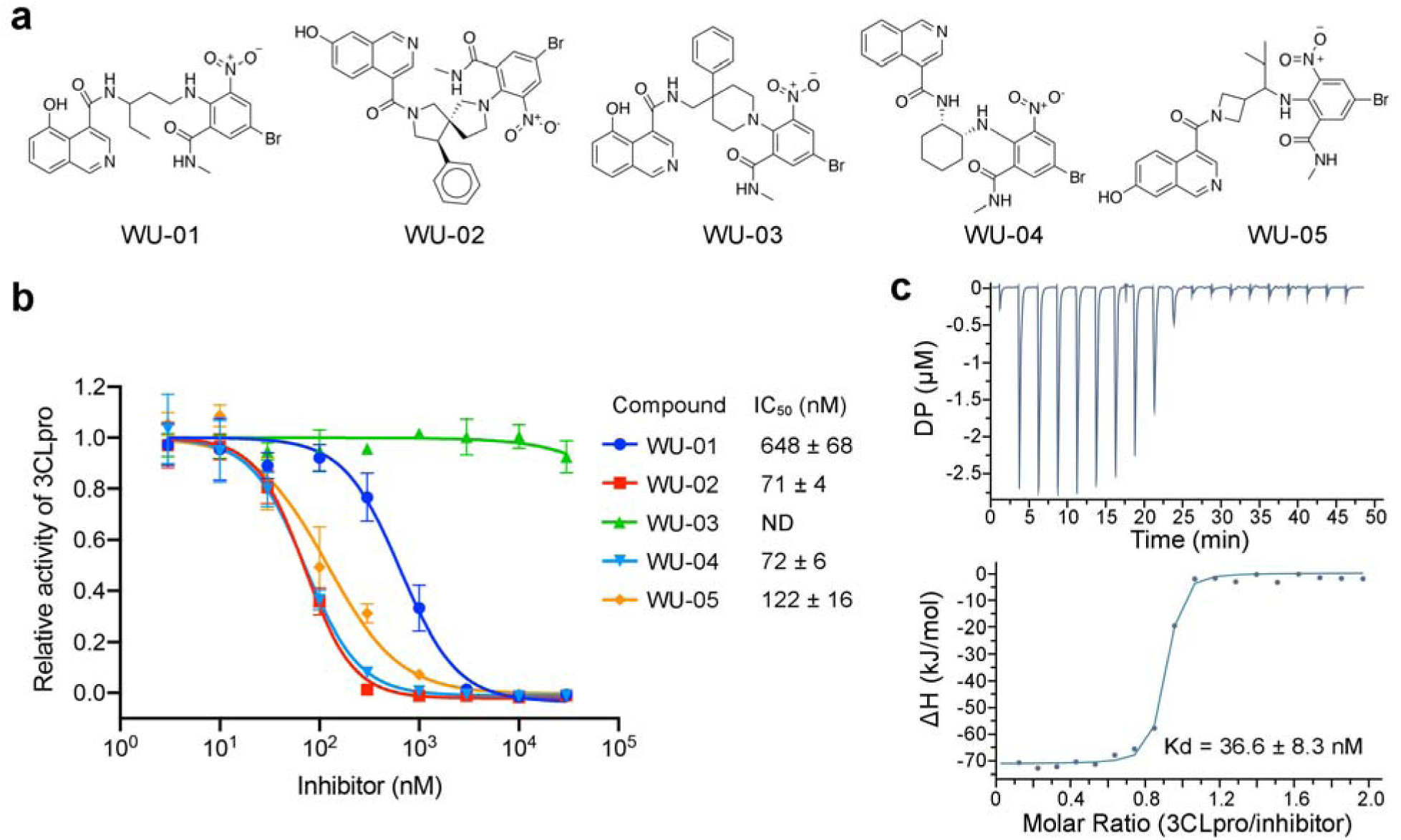
Identification of non-covalent inhibitors of SARS-CoV-2 3CLpro. **a**, Five compounds containing an isoquinoline ring and a bromophenyl ring were identified as 3CLpro binders. **b**, The ability of the 3CLpro binders to inhibit the enzyme activity of SARS-CoV-2 3CLpro was evaluated using the fluorescent substrate Dabcyl-KTSAVLQSGFRKME-Edans. WU-02 and WU-04 showed the highest inhibitory activity. The data represents the mean ± SD of three independent measurements. **c**, The binding affinity (*K*_d_) between WU-04 and SARS-CoV-2 3CLpro was measured using isothermal titration calorimetry (ITC).

We also evaluated the ability of WU-04 to inhibit the 3CLpro of the SARS-CoV-2 Omicron variant, which has a single mutation P132H. WU-04 inhibited the 3CLpro P132H mutant with an IC_50_ of 53 nM, similar to that against the wild-type 3CLpro (fig. S2).

### Mechanism of inhibition by non-covalent inhibitors

To understand how these compounds bind to and inhibit 3CLpro, we co-crystallized the SARS-CoV-2 3CLpro with compounds WU-02 and WU-04. The structures were determined using molecular replacement and refined to resolutions of 1.90 Å and 1.83 Å, respectively (Table S1). Both WU-02 and WU-04 bind to 3CLpro with a 1:1 ratio and in a similar pose (Fig. 2a and fig. S3, S4a and S4b). The following structure description will focus on our frontrunner WU-04. The structure of 3CLpro/WU-02 is detailed in figures S2 and S3b.

**Fig. 2.**
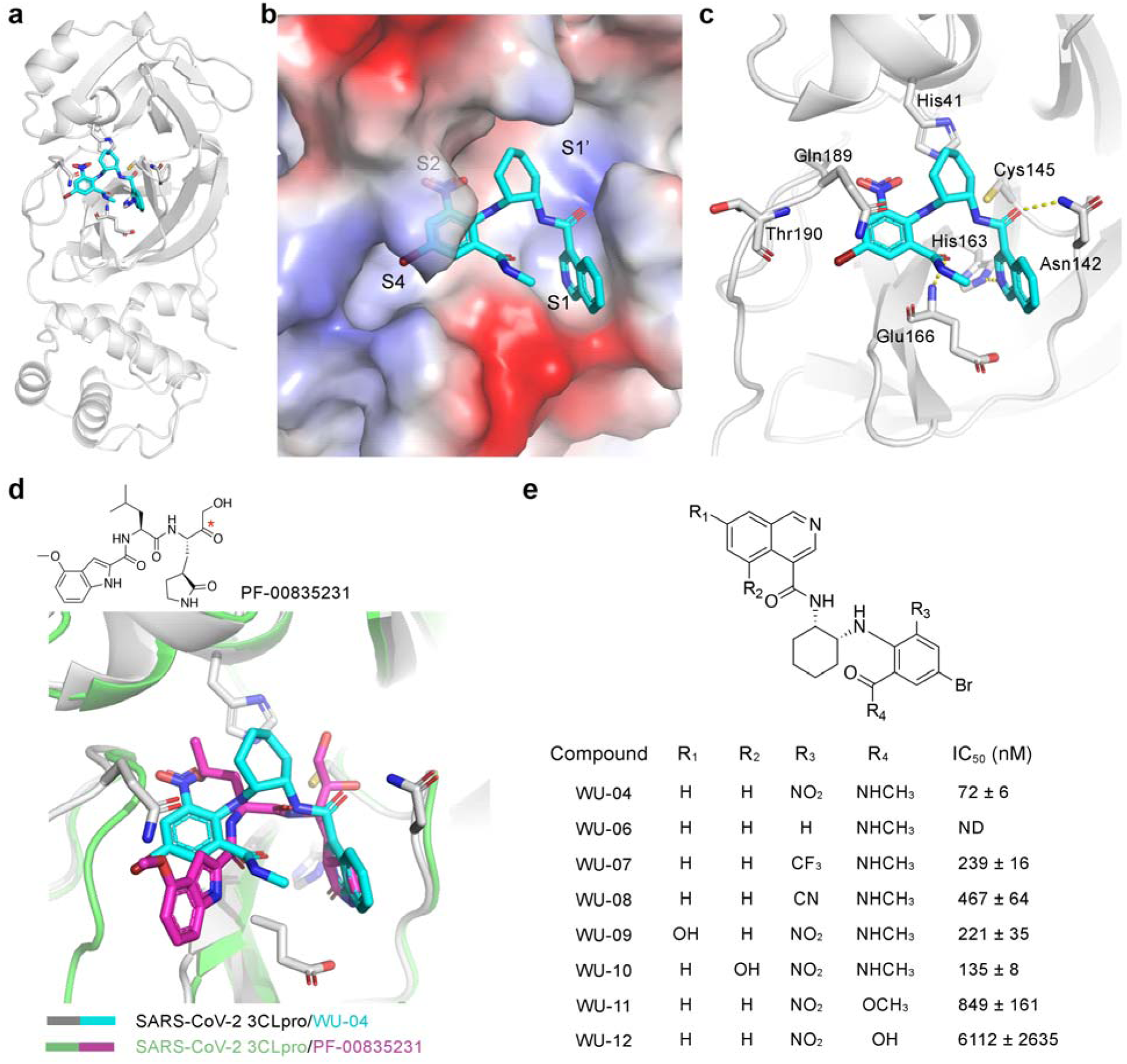
The non-covalent inhibitor WU-04 binds into the catalytic pocket of SARS-CoV-2 3CLpro. **a**, The overall structure of the 3CLpro/WU-04 complex. **b**, WU-04 (cyan) binds into the catalytic pocket of 3CLpro. The catalytic pocket consists of S1’, S1, S2 and S4 sites. The protein contact potential was calculated using PyMOL. **c**, Details of the interaction between WU-04 (cyan) and residues around the catalytic pocket of 3CLpro. Hydrogen bonds are indicated by yellow dash lines. **d**, Alignment of the crystal structure of the SARS-CoV-2 3CLpro/WU-04 complex with that of the SARS-CoV-2 3CLpro/PF-00835231 complex (PDB ID: 6XHM). The reaction site of PF-00835231 is indicated by a red asterisk. **e**, Seven analogs of WU-04 were synthesized and their inhibitory activity against SARS-CoV-2 3CLpro was evaluated using the fluorescent substrate Dabcyl-KTSAVLQSGFRKME-Edans. The data represents the mean ± SD of three independent measurements.

In the crystal structure, WU-04 binds to the catalytic pocket of 3CLpro, indicating that WU-04 blocks the access of 3CLpro substrates to the catalytic dyad (Fig. 2b and 2c, and fig. S3a). WU-04 is thus a competitive inhibitor of 3CLpro. According to the interactions between 3CLpro and its peptide substrates, the catalytic pocket of 3CLpro can be divided into four sites: S1’, S1, S2 and S4 (36). The isoquinoline ring of WU-04 docks into the S1 site. The nitrogen atom of the isoquinoline ring and the carbonyl group linked to the isoquinoline ring form hydrogen bonds with the side chains of His163 and Asn142, respectively (Fig. 2c). The 6-nitro and 4-bromo groups of the bromophenyl ring occupy sites S2 and S4, respectively (Fig. 2b). The amino-π interaction between the phenyl ring and the side chain of Gln189 contributes to the potency of WU-04 (Fig. 2c). The strong electron-withdrawing ability of the nitro group makes the aromatic ring positively charged, thereby enhancing this amino-π interaction (37). Such electron-withdrawing ability also enhances the strength of the halogen bond with the carbonyl group of Thr190. In addition, the carbonyl oxygen of the methylcarbamoyl group of WU-04 accepts a hydrogen bond from the main chain amide of Glu166 (Fig. 2c).

We compared the binding mode of WU-04 with six representative covalent inhibitors and three non-covalent inhibitors of SARS-CoV-2 3CLpro (Fig. 2d and fig. S5). PF-00835231 is a covalent inhibitor developed by Pfizer to target the 3CLpro of SARS-CoV and SARS-CoV-2 (14). In PF-00835231, a γ-lactam moiety docks into site S1, mimicking the side chain of glutamine at the P1 position of the substrates of 3CLpro. Similar to the isoquinoline ring in WU-04, the carbonyl oxygen of the γ-lactam moiety forms a hydrogen bond with His163 of 3CLpro (14). Thus, a moiety that can accept a hydrogen bond from His163 is important for the binding of inhibitors to 3CLpro. In other reported structures, a γ-lactam (fig. S5b, 5c, 5d and 5f), a cyclobutyl (fig. S5e), a pyridine (fig. S5g), a benzotriazol (fig. S5h), or a 1-methyl-1H-1,2,4-triazol moiety inserts into site S1 of 3CLpro (11,12,15,16,24,27,30,31). The major difference between WU-04 and the reported inhibitors is the presence of the bromophenyl ring in WU-04 that fits well into sites S2 and S4. As we mentioned above, the bromophenyl ring interacts with Gln189 and Thr190 through amino-π interaction and halogen bonding, respectively (Fig. 2c and fig. S5a). Incorporating the bromophenyl ring moiety into the covalent inhibitors is expected to increase their potency.

Based on structural insights and free energy perturbation calculations, we designed and synthesized a series of WU-04 analogs to explore its structure-activity relationship (Fig. 2e). Replacement of the nitro group with hydrogen (WU-06) abolished the activity of WU-04, indicating that the nitro group is essential. Changing the nitro group to other electron-withdrawing groups, such as trifluoromethyl and cyano groups (WU-07 and WU-08), reduced the activity. Adding a hydroxyl group to the 7- or 5-position of the isoquinoline ring (WU-09 and WU-10) slightly decreased the activity. Changing the methylcarbamoyl group to a carboxyl methyl ester (WU-11) decreased the potency by an order of magnitude, while changing the same group to a carboxyl group (WU-12) further decreased the inhibition. The methylcarbamoyl group is in the vicinity of Glu166 in 3CLpro. Incorporating a negatively charged moiety such as a carboxyl group at this position might introduce unfavorable electrostatic interactions with Glu166, thereby destabilizing the binding of the inhibitor to 3CLpro.

### 3CLpro inhibitors inhibit SARS-CoV-2 replication in cellular assays

We next tested whether WU-02 and WU-04 could block SARS-CoV-2 replication in cellular assays. The antiviral activities were evaluated using a luciferase SARS-CoV-2 infecting a human alveolar epithelial cell line that overexpresses the human angiotensin-converting enzyme 2 (A549-hACE2) (38). Both WU-02 and WU-04 potently inhibited the replication of SARS-CoV-2, with EC_50_ values of 54 and 12 nM, respectively, and EC_90_ values of 305 and 36 nM, respectively (Fig. 3a). A three-day toxicity of the two compounds in A549 cells was also evaluated (Fig. 3b). Both exhibited 50% cytotoxic concentration (CC_50_) values greater than 20 µM (the highest compound concentration used in the assay). We also evaluated both compounds’ anti-SARS-CoV-2 activities using SARS-CoV-2 with a firefly luciferase reporter in human lung adenocarcinoma epithelial Calu-3 cells. The EC_50_ values of WU-02 and WU-04 were 80 and 25 nM, respectively (Fig. 3c), similar to those in A549-hACE2 cells. WU-04 also exhibited a high potency against SARS-CoV-2 in primary normal human bronchial epithelial (NHBE) cells, with an EC_50_ of around 3 nM (fig. S6a), as well as in Vero E6 cells, with an EC_50_ of around 10 nM (fig. S6b).

**Fig. 3.**
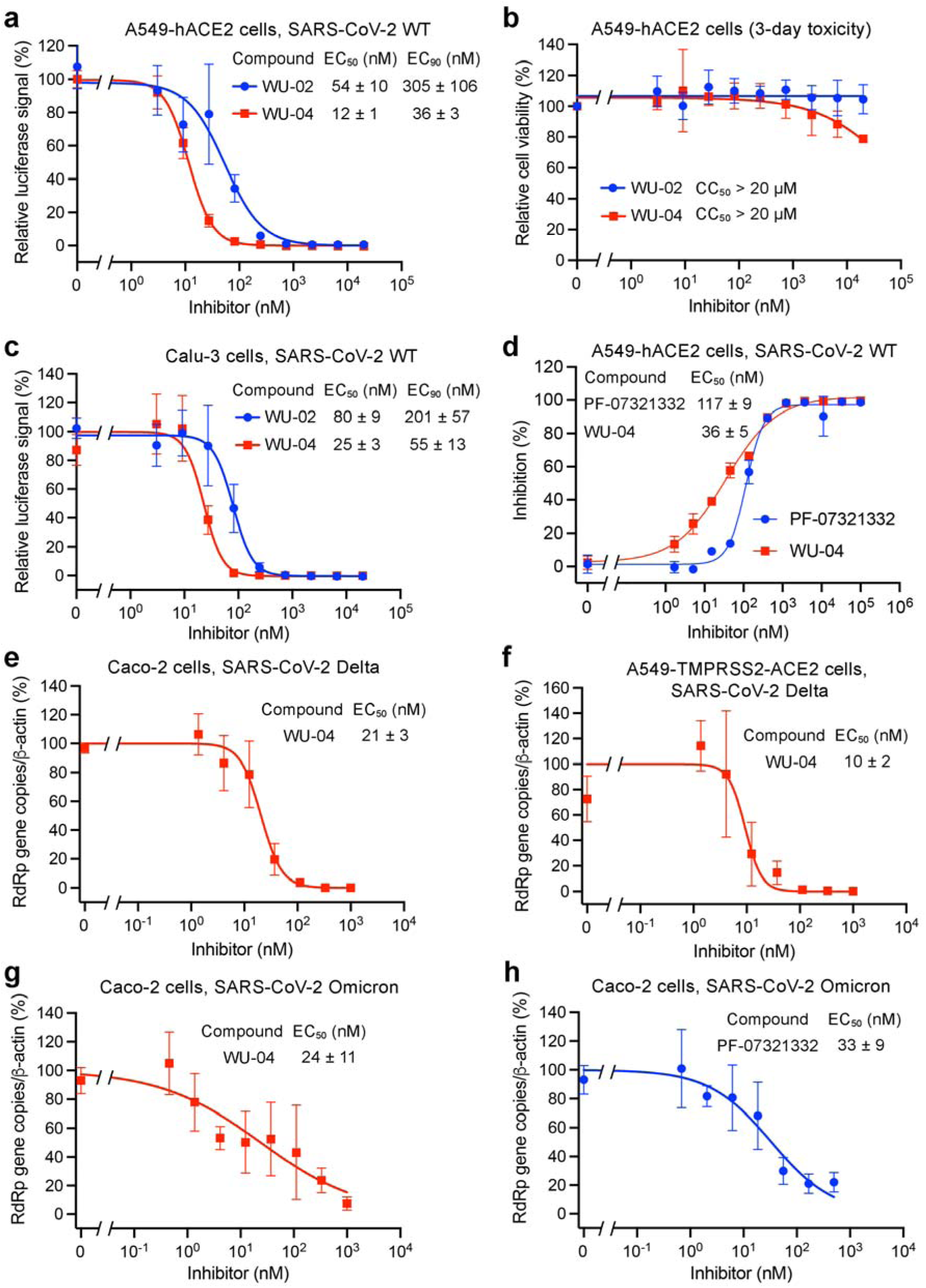
Inhibition of SARS-CoV-2 replication by 3CLpro inhibitors in human cell lines. **a** and **c**, The anti-SARS-CoV-2 (wild-type strain) activity of WU-02 and WU-04 was measured in A549-hACE2 cells (**a**), or in Calu-3 cells (**c**) using a nanoluciferase SARS-CoV-2 assay (38). **b**, Three-day cytotoxicity of WU-02 and WU-04 was evaluated in A549-hACE2 cells. **d**, The anti-SARS-CoV-2 (wild-type strain) activity of WU-04 and that of PF-07321332 were evaluated in A549-hACE2 cells by quantifying the expression of the viral nucleocapsid protein using immunofluorescence microscopy. **e** and **f**, The anti-SARS-CoV-2 (Delta variant) activity of WU-04 was measured in Caco-2 cells (**e**) or A549-TMPRSS2-ACE2 cells (**f**) by using RT-qPCR to determine the viral RdRp gene copy number. **g** and **h**, The anti-SARS-CoV-2 (Omicron variant) activity of WU-04 (**g**) and that of PF-07321332 (**h**) were measured in Caco-2 cells by using RT-qPCR to determine the viral RdRp gene copy number. The data represent the mean ± SD of four to eight (**a** to **c**), two (**d**), or three (**e** to **h**) independent measurements.

The anti-SARS-CoV-2 activity of WU-02 was also evaluated in A549-hACE2 cells by quantifying the expression of the viral nucleocapsid (N) protein (39), and it was compared with the activity of Pfizer’s compound PF-07321332. The EC_50_ of WU-04 was around 36 nM, better than that of PF-07321332 (EC_50_ = 117 nM) (Fig. 3d).

Several SARS-CoV-2 variants have emerged. We evaluated the antiviral activity of WU-04 against the Delta variant and the Omicron variant. In Caco-2 cells and in A549 cells overexpressing human TMPRSS2 and ACE2 (A549-TMPRSS2-ACE2), the EC_50_ of WU-04 against the Delta variant were 21 nM and 10 nM, respectively (Fig. 3e and 3f). In Caco-2 cells, the EC_50_ of WU-04 against the Omicron variant was around 24 nM, slightly better than that of PF-07321332 (EC_50_ = 33 nM) (Fig. 3g and 3h).

### WU-04 is a pan-inhibitor of coronavirus 3CLpro

In addition to the anti-SARS-CoV-2 activity, the inhibitory activity of WU-04 against 3CLpro of SARS-CoV and MERS-CoV was also evaluated. WU-06 was used as the negative control.

In a fluorescence-based enzymatic activity assay in which the concentrations of 3CLpro and the fluorogenic substrate were 100 nM and 100 µM, respectively, WU-04 potently inhibited the SARS-CoV 3CLpro, with an IC_50_ value of 55 nM, while WU-06 showed no activity (Fig. 4a). The binding affinity between WU-04 and SARS-CoV 3CLpro was about 65 nM (fig. S7a). We also determined the crystal structure of SARS-CoV 3CLpro bound to WU-04 at a resolution of 1.99 Å (Table S1, Fig. 4b and fig. S4c). Consistent with the enzymatic assay results and the binding affinity, the key residues of the inhibitor-binding pocket of SARS-CoV 3CLpro and the interactions between these residues and WU-04 are almost identical to those observed in the SARS-CoV-2 3CLpro/WU-04 structure (Fig. 2c). The anti-SARS-CoV activity of WU-04 was measured in Calu-3 cells (Fig. 4c) and in Vero E6 cells (fig. S7c), and the EC_50_s were around 10-19 nM.

**Fig. 4.**
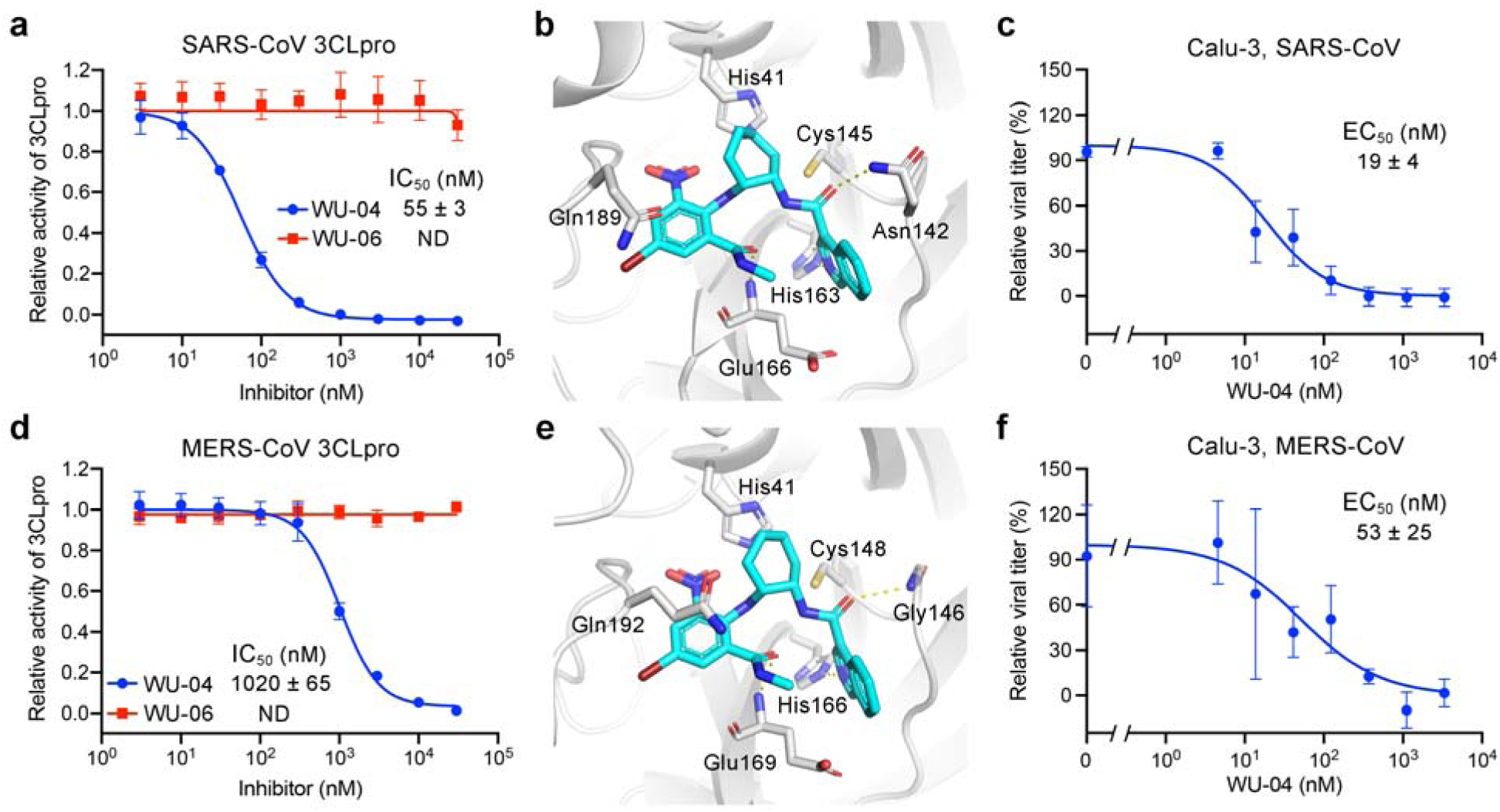
Compound WU-04 also inhibited the 3CLpro of SARS-CoV and MERS-CoV. **a** and **d**, WU-04 but not WU-06 inhibited the enzyme activity of SARS-CoV 3CLpro (**a**) and MERS-CoV 3CLpro (**d**). The data represent the mean ± SD of three independent measurements. **b** and **e**, Close-up view of the binding modes of WU-04 to SARS-CoV 3CLpro (**b**) and MERS-CoV 3CLpro (**e**). WU-04 is colored cyan. Hydrogen bonds are represented by yellow dash lines. **c** and **f**, The antiviral activity of WU-04 against SARS-CoV (**c**) or against MERS-CoV (**f**) was measured in Calu-3 cells. The data represents the mean ± SD of three independent measurements.

The 3CLpro of MERS-CoV shares only 50% sequence identity with SARS-CoV-2 3CLpro. Even so, WU-04 inhibited MERS-CoV 3CLpro with an IC_50_ of about 1 µM (Fig. 4d). The concentrations of MERS-CoV 3CLpro and the fluorogenic substrate used in the assay were 500 nM and 200 µM, respectively, which were higher than those used in the SARS-CoV-2 and SARS-CoV 3CLpro assays, because MERS-CoV 3CLpro is a weakly associated dimer in contrast to SARS-CoV-2 3CLpro and SARS-CoV 3CLpro (40), and thus a higher concentration is required to facilitate the dimerization. The higher concentrations of the MERS-CoV 3CLpro and its substrate inflated the IC_50_ value of WU-04 against this 3CLpro. Consistent with WU-04 being an effective inhibitor of MERS-CoV 3CLpro, its binding affinity (*K*_d_) to MERS-CoV 3CLpro was about 32 nM (fig. S7b). The crystal structure of the MERS-CoV 3CLpro/WU-04 complex was also determined using molecular replacement and refined to 2.98 Å resolution (Table S1, Fig 4e and fig. S4d). The binding mode of WU-04 in this structure was very similar to that in the structure of the SARS-CoV-2 3CLpro/WU-04 complex (Fig. 4e). The anti-MERS-CoV activity of WU-04 was confirmed in Calu-3 cells and in Vero E6 cells, giving the EC_50_ values of around 53 nM and 609 nM, respectively (Fig. 4f and fig. S7d).

Alignment of the 3CLpro sequence of SARS-CoV-2 with those of SARS-CoV, MERS-CoV and other four human coronaviruses shows that most key residues in direct contact with WU-04 are conserved; in addition, most other residues within 5 Å of WU-04 in the SARS-CoV-2 3CLpro structure are also conserved (fig. S8). We conclude that WU-04 is a pan-inhibitor of coronavirus 3CLpro.

### WU-04 inhibits SARS-CoV-2 replication in mice models

The ability of WU-04 to inhibit SARS-CoV-2 replication was evaluated in two mouse models. The first was BALB/c mice infected with a mouse-adapted SARA-CoV-2 variant (41). The mice were given oral administration of 250 mpk (mg/kg of body weight per dose) of WU-04 (twice daily) or the vehicle control; the first dose was administered one hour before the infection (Fig. 5a). At day 3 post-infection, the mice were euthanized, and the nasal turbinate and lung tissues were harvested. Upon treatment with WU-04, the viral RNA copy numbers and viral titers in the nasal turbinates of three of the six mice, and that in the lungs of five of the six mice, were decreased to the detection limit; in comparison with the vehicle group, the average viral RNA copy number and the average viral titer of the WU-04 group had a reduction of 5 and 3 log units, respectively (Fig. 5b and 5c).

**Fig. 5.**
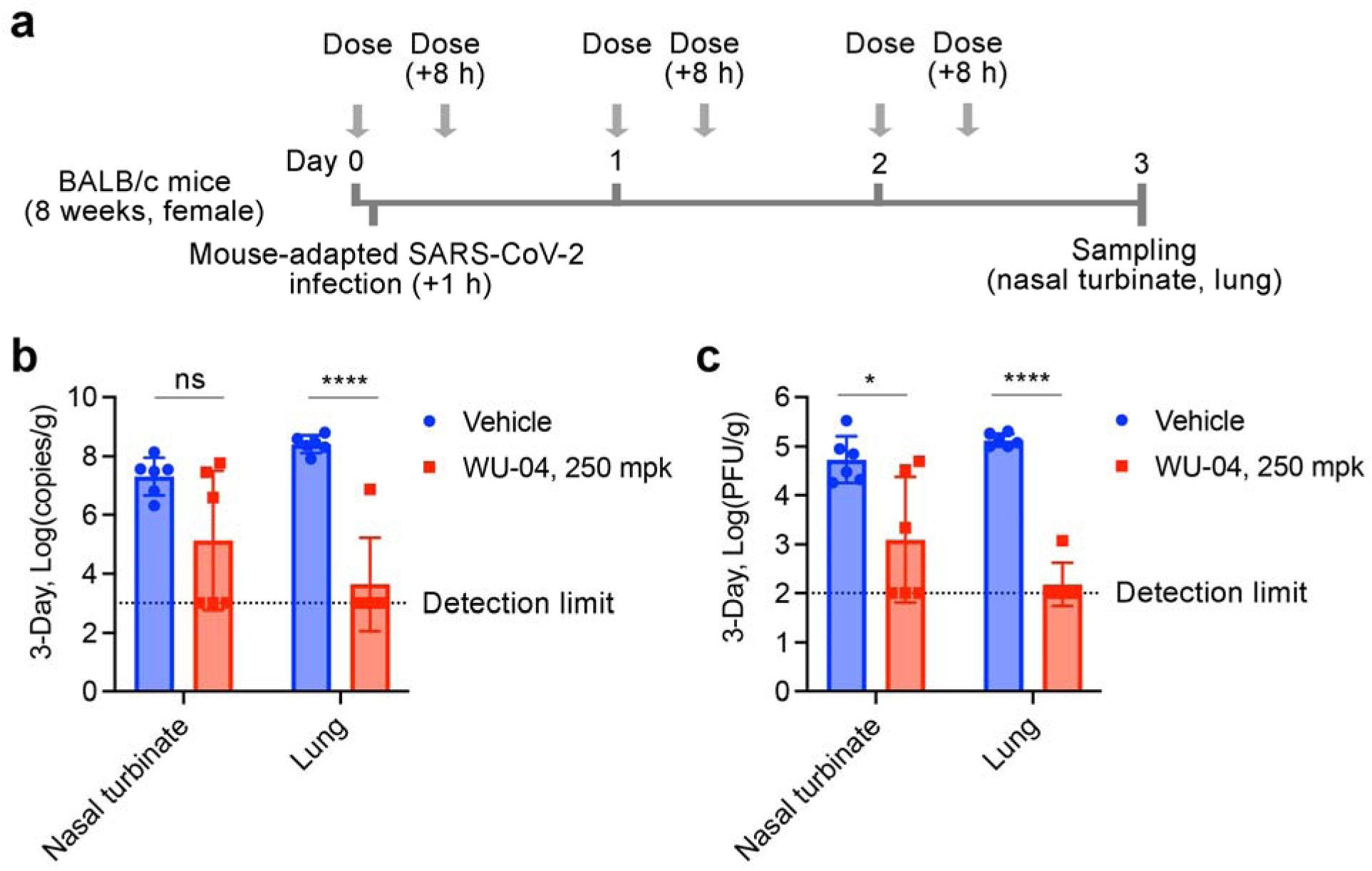
In vivo efficacy of WU-04 against a mouse-adapted SARS-CoV-2 variant in mice. **a**, Schematic diagram of the study process. **b** and **c**, After three-day treatment with WU-04 or the vehicle control, the viral RNA copy numbers (**b**) and the viral titers (**c**) in the nasal turbinate and lung of each mouse were determined. There were six mice in each group. * P < 0.05, **** P < 0.0001.

The second model was K18-hACE2 transgenic mice, for which SARS-CoV-2 infection can cause severe lung inflammation and death (42,43). After infection, the mice were orally administered with WU-04 (100, 200, or 300 mpk, twice daily), or PF-07321332 (300 mpk, twice daily), or the vehicle control (Fig. 6a). The average body weight of the mice in the vehicle group began to decrease at two days postinfection and decreased about 5% one day later; in contrast, the average body weight in the inhibitor-treated groups did not change (300 mpk of PF-07321332, 100 mpk of WU-04) or slightly increased (200 and 300 mpk of WU-04) (Fig. 6b). WU-04 showed a concentration-dependent anti-SARS-CoV-2 activity. With a dose of 100 mpk, WU-04 slightly decreased the viral RNA levels in the nasal turbinates and brains (Fig. 6c). With a dose of 200 mpk, WU-04 reduced the viral RNA by more than 1.5 log units in the nasal turninates, lungs and brains of the infected mice compared with the vehicle group (Fig. 6c). With a dose of 300 mpk, the viral RNA in the nasal turbinates and lungs of four of the six WU-04 treated mice decreased to the detection limit, similarly to the PF-07321332 treated group; in brains, WU-04 and PF-07321332 also showed similar activities to reduce the viral RNA (Fig. 6c). Measurement of the viral titers also supported the concentration-dependent anti-SARS-CoV-2 activity of WU-04 (Fig. 6d). Histopathology analysis of the lung tissues and immunohistochemical analysis of the brain tissues showed that WU-04 with a dose of 300 mpk protected the lung from SARS-CoV-2 induced inflammation (Fig. 6e and 6f), and blocked SARS-CoV-2 infection in brain (Fig. 6g). These results validate the *in vivo* efficacy of WU-04 and demonstrate that WU-04 had an antiviral activity similar to that of PF-07321332 when the same dose was orally administered.

**Fig. 6.**
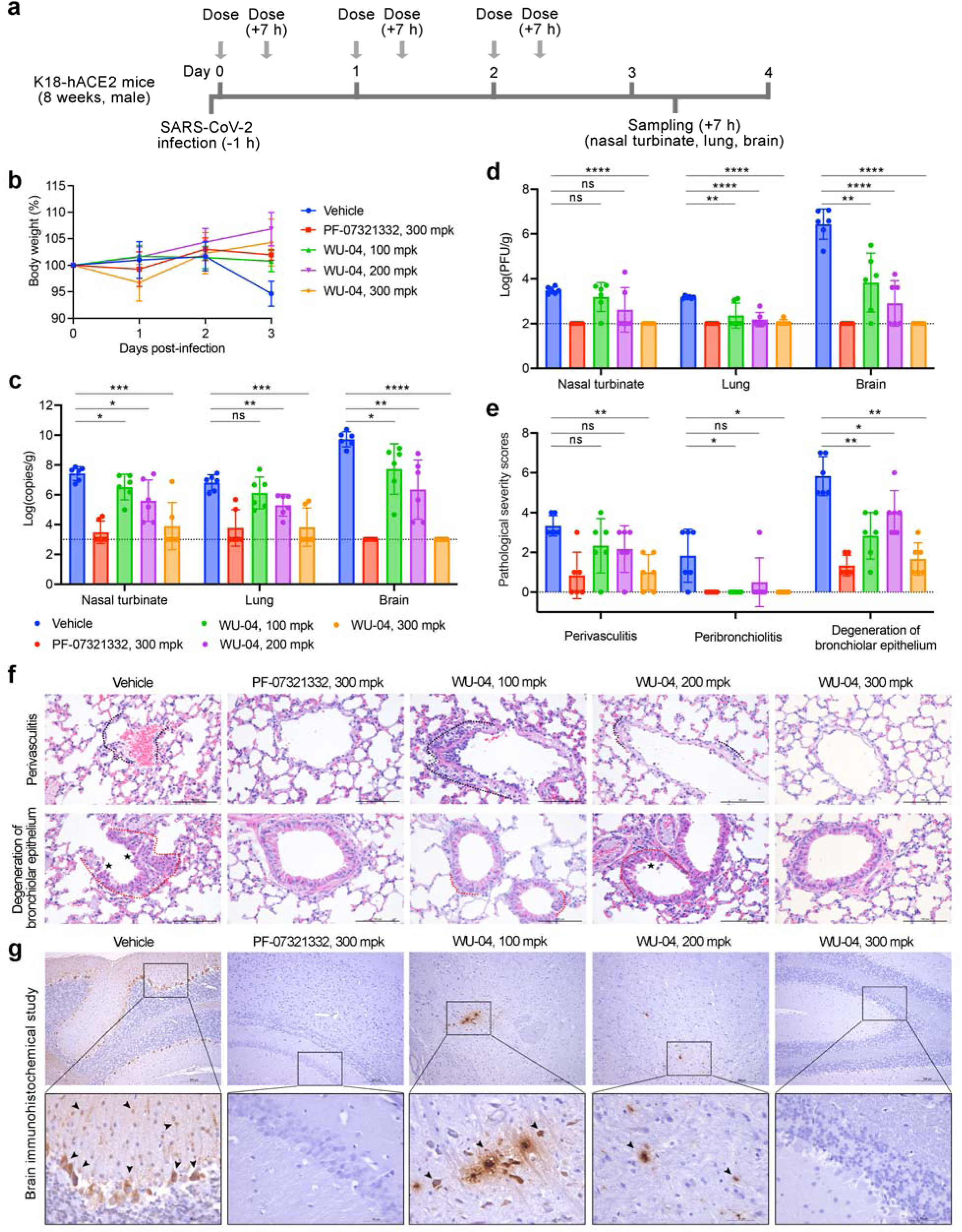
In vivo efficacy of WU-04 against a SARS-CoV-2 in K18-hACE2 mice. **a**, Schematic diagram of the study process. **b**, Changes of the mouse body weights during the study. **c** and **d**, After three days treatment with WU-04, PF-07321332, or the vehicle control, the viral RNA copy numbers (**c**) and the viral titers (**d**) in the nasal turbinate, lung and brain of each mouse were determined. **e, f** and **g**, After three days treatment, the lung tissues were harvested and processed for histological analysis (**f**) and the pathological changes were scored (**e**), and the brain tissues were harvested and processed for immunohistochemical analysis (**g**) using methods in the supplementary materials. Severe lymphoplasmacytic perivasculitis and vasculitis (black dotted line), peribronchiolar inflammatory infiltrate (red dotted line), degeneration of bronchiolar epithelium (asterisk), viral antigen-positive cells in the brain (arrow) were marked. * P < 0.05, ** P < 0.01, *** P < 0.001, ***** P < 0.0001.

### Pharmacokinetics of WU-04

We evaluated the pharmacokinetics of WU-04 through *in vitro* and *in vivo* studies. To examine which isoforms of cytochromes P450 (CYP) are responsible for the metabolism of WU-04, we measured the inhibition of WU-04 metabolism in human liver microsomes by CYP isoform-selective inhibitors. The CYP2C8 inhibitor montelukast and the CYP3A inhibitor ketoconazole inhibited 9.8% and 84.4% of the metabolism of WU-04, respectively; the inhibitors of other CYP isoforms had little effect on the metabolism of WU-04 (Table S2). We conclude that CYP3A subfamily members are the major CYP isoforms involved in WU-04 metabolism. Next, we tested the effect of ritonavir (RTV) on WU-04 metabolism in liver microsomes. RTV is a CYP3A4 inhibitor and has been used together with PF-07321332 to treat COVID-19 patients (44). Without RTV, the half-life of WU-04 in human, mouse, and dog liver microsomes were between 0.9 to 2.3 minutes; in contrast, in the presence of RTV, the half-life was increased to more than 145 minutes in human liver microsomes, and 87.1 minutes and 69.7 minutes in mouse and dog liver microsomes, respectively (Table S3).

In *in vivo* studies, we first evaluated the pharmacokinetics of WU-04 in BALB/c mice (fig. S9a to S9d). After oral administration, the half-life of WU-04 in the plasma were 0.25 hour, 1.49 hours, and 2.02 hours at a dose of 50 mpk, 125 mpk, and 300 mpk, respectively, in a dose-dependent manner; when WU-04 was administered together with 20 mpk of RTV, the half-life was increased to 1.54 to 4 hours. RTV significantly increased the exposure (AUC, area under the plasma concentration-time curve) of WU-04: at a dose of 50 mpk, the AUC of WU-04 was increased from 1260 to 51051 ng*hour/mL; at a dose of 300 mpk, the AUC was increased from 69796 to 488615 ng*hour/mL (fig. S9d).

In Beagle dogs, the half-life of WU-04 in the plasma was 1.90 hours when 1 mpk of WU-04 was administered via intravenous (IV) bolus route, and was 1.68 hours when 5 mpk of WU-04 was administered via oral route (PO) (fig. S9e and S9g). The calculated bioavailability of WU-04 was about 18%. To evaluate the effect of RTV on the pharmacokinetics of WU-04 in dogs, 3.5 mpk of RTV was administered 12 hours before 5 mpk of WU-04 was administered together with 3.5 mpk of RTV, 12 hours later, a third dose of RTV (3.5 mpk) was administered; RTV increased the half-life of WU-04 (5 mpk) to 4.83 hours and increased the exposure of WU-04 from 3670 to 38494 ng*hour/mL (fig. S9f and S9g). When WU-04 was administered orally with a dose of 100 mpk, the plasma concentration-time curve was similar to that when 5 mpk of WU-04 was administered together with 3.5 mpk of RTV (fig. S9f).

The unbound ratio of WU-04 in CD-1 mouse plasma, Beagle dog plasma, and in human plasma were 2.3%, 4.2%, and 5.2%, respectively. According to the EC_90_ values of WU-04 in figures 3a and 3c (36 nM and 55 nM, respectively), the total EC_90_ of WU-04 in mice were 824 ng/mL and 1259 ng/mL, respectively, while in dogs were 451 ng/mL and 689 ng/mL, respectively. We found that at a dose of 300 mpk, the plasma concentration of WU-04 in mice fell below the total EC_90_ values 5-6 hours after dosing (fig. S9c); meanwhile, 300 mpk of WU-04 showed antiviral activity similar to that of 300 mpk of PF-07321332 (Fig. 6). In Beagle dogs, when 5 mpk of WU-04 was administered together with 3.5 mpk of RTV, the plasma concentration of WU-04 at 12 hours post dosing was 597 ng/mL, close to the total EC_90_ values (fig. S9f). The pharmacokinetics studies of WU-04 in human have not been carried out yet; but considering that with the help of RTV, WU-04 had a longer half-life in human liver microsomes than in dog liver microsomes (Table S3), and the unbound ratio of WU-04 in human plasma was higher than in dog plasma, we speculate that in combination with RTV, WU-04 with a dose of no more than 5 mpk (twice daily) may be effective to treat COVID-19 patients.

## Discussion

Since the outbreak of COVID-19, several small molecule inhibitors of SARS-CoV-2 have been reported, but only three have been approved for the treatment of COVID-19, including remdesivir, molnupiravir, and nirmatrelvir (PF-07321332). PF-07321332 inhibits the viral protease 3CLpro while remdesivir and molnupiravir block the viral RNA replication. PF-07321332, in combination with ritonavir, can be administered orally and showed better efficacy in COVID-19 clinical trial (44). Some other 3CLpro inhibitors with *in vivo* anti-coronavirus efficacy have been reported: most of them share similar scaffold as PF-07321332 and use an electrophile to covalently link to the catalytic cysteine of 3CLpro (12,19,21,22,45), and two of them are non-covalent inhibitors (31,46). Even though, SARS-CoV-2 inhibitors with higher potency, lower toxicity, and good bioavailability are still needed. We have identified WU-04 as a potent non-covalent inhibitor of SARS-CoV-2 3CLpro. WU-04 also bound to the 3CLpro of SARS-CoV and MERS-CoV with high affinity and inhibited their enzyme activity. WU-04 showed high potency in all cell lines tested. In A549 cells, it had higher antiviral activity than PF-07321332. In addition, the antiviral activity comparable to PF-07321332 in K18-hACE2 mice, and the pharmacokinetics of WU-04 indicate that WU-04 is a promising drug candidate for the treatment of COVID-19 and other coronavirus diseases. The novel scaffold of WU-04 and the cocrystal structures of SRAS-CoV, SARS-CoV-2, and MERS-CoV 3CLpro in complex with WU-04 enable rational design of the current inhibitors and future drug discovery.

## Acknowledgments

We thank Dr. Sheng-ce Tao at Shanghai Jiao Tong University for sharing the plasmids of SARS-CoV-2 proteins. We thank the Protein Characterization and Crystallography Facility of Westlake University for assistance in crystallization and X-ray data collection. We thank the Instrumentation and Service Center for Molecular Sciences at Westlake University for supporting in 1H-NMR and LC-MS data measurement. We would like to express our sincere gratitude towards Tencent Foundation for their generous support in this collaborative project.

## Funding

This work was supported by:

Westlake Education Foundation (J.H., Q.H.)

Tencent Foundation (J.H., Q.H.)

National Natural Science Foundation of China (21803057, 32041005)

Zhejiang Provincial Natural Science Foundation of China (LR19B030001)

The National Key Research and Development Program of China (2021YFC2301700, 2020YFA0707701)

## Author contributions

H.Y., P.Y.S., J.H., and Q.H. conceived the project. N.H., L.Z., and C.P. purified the proteins and performed the biochemical assays. N.H., C.P. and H.G. performed the crystallization experiments. L.Z. synthesized compounds WU-09 to WU-12. X.X., K.T., Y.Z., Y.Y., J.Z., C.S., and V.M. designed and performed the cell-based antiviral assays. L.S., G.Z., Z.W., C.W., X.H., H.H., performed the antiviral assays in mouse models. W.Z., Q.T., Y.X., and J.X. performed the computational docking and free energy calculations. Q.H. designed the DEL selection conditions. W.S., Y.Y., and Z.S. performed the DEL selection. N.H., L.S., L.Z, X.X., R.Z., S.Y., Z. B., H.Y., P.Y.S., J.H., and Q.H. analyzed the data. H.Y., P.Y.S., J.H., and Q.H. wrote the manuscript.

## Data and materials availability

The crystal structures have been deposited in the Protein Data Bank (www.rcsb.org) with the accession numbers 7EN9 (SARS-CoV-2 3CLpro/WU-02 complex), 7EN8 (SARS-CoV-2 3CLpro/WU-04 complex), 7END (SARS-CoV 3CLpro/WU-04 complex) and 7ENE (MERS-CoV 3CLpro/WU-04 complex). All other data are available in the manuscript or the supplementary materials.

## Supplementary Materials

Materials and Methods

Figs. S1 to S9

Tables S1 to S3

