## Supplementary Materials for "Development of highly potent non-covalent inhibitors of SARS-CoV-2 3CLpro"

**This PDF file includes:**

Materials and Methods
Figs. S1 to S9
Tables S1 to S3

### Materials and Methods

#### Biological experiments

Genes and cloning

The gene coding for the SARS-CoV-2 3CLpro was a gift from Prof. Sheng-ce Tao at Shanghai Jiao Tong University. The genes of SARS-CoV 3CLpro and MERS-CoV 3CLpro were optimized for *Escherichia coli* expression and synthesized at GENEWIZ, Suzhou, China. The expression plasmids of SARS-CoV-2 3CLpro and SARS-CoV 3CLpro were constructed by following a protocol published previously (1). The expression plasmid of MERS-CoV 3CLpro was constructed by introducing the full-length gene of MERS-CoV 3CLpro with a Factor Xa cleavage site at its N terminus and a stop codon at its C terminus into pET15b vector.

Protein expression and purification

The 3CLpro of SARS-CoV-2 and SARS-CoV were purified following a method described previously (1). The expression plasmid was transformed into *Escherichia coli* BL21(DE3). The cells were cultured in LB medium containing 0.1 mg/ml ampicillin at 37°C until the absorbance at 600 nm (OD600) reached 0.6, then 0.5 mM β-D-thiogalactopyranoside (IPTG) was added and the cell culture was incubated at 16 °C for 12 hours. As for MERS-CoV 3CLpro, 0.3 mM IPTG was added, and the cell culture was incubated at 22 °C for 12 hours. The cells were harvested by centrifugation at 4,000 g for 10 minutes (Thermo Fisher LYNX 6000 Centrifuge, F9-6x 1000). The cells were resuspended in lysis buffer (40 mM Tris-HCl, pH 8.0, 100 mM NaCl) supplemented with 1 mM phenylmethylsulfonyl fluoride (PMSF), and lysed by ultrasonication (SONICS, vibra cell). The cell lysate was centrifuged at 30,000 g (BECKMAN Avanti JXN-26 Centrifuge, JA-14.50) for 45 minutes. The supernatant was incubated with $Co^{2+}$ resin (TALON, Cat# 635504) at

4 °C for 1 hour, then the resin was transferred into gravity columns and washed with the wash buffer (40 mM Tris-HCl pH 8.0, 500 mM NaCl, 5 mM imidazole). 3CLpro was eluted from the resin by the elution buffer (40 mM Tris-HCl, pH 8.0, 50 mM NaCl, 250 mM imidazole). After adding 5 mM dithiothreitol (DTT) and 10% glycerol, the eluate was incubated with human rhinovirus 3C protease (TaKaRa, Cat# 7360) (for the 3CLpro of SARS-CoV-2 and SARS-CoV) or with Factor Xa protease (NEB, Cat# P8010) at 4 °C for 12 hours to remove the 6xHis tag. The efficiency of cleavage was analyzed by SDS-PAGE. After cleavage, the sample was first diluted with buffer A (25 mM Tris-HCl, pH 8.0) to decrease the NaCl concentration to about 5 mM, then loaded into a Source-15Q column (GE Healthcare) and eluted by a linear gradient from 100% buffer A (25 mM Tris 8.0) to 40% Buffer B (25 mM Tris 8.0, 1 M NaCl). The purity of each fraction was analyzed by SDS-PAGE. If there is still 3CLpro with a 6xHis tag in the sample, it can be removed by loading the sample onto a $Co^{2+}$ resin column (TALON, Cat# 635504). The 3CLpro was further purified by gel filtration. Briefly, the protein was supplemented with 5 mM DTT, concentrated, loaded into a Superdex 200 increase 10/300 GL column (GE Healthcare), and then eluted with 20 mM HEPES, pH 7.4, 150 mM NaCl. The peak fractions were pooled and concentrated to about 10 mg/mL and stored at -80 °C for subsequent biological assays and crystallization.

Affinity selection against 6xHis-tagged 3CLpro

HisPur™ Ni-NTA Magnetic Beads (Thermo Scientific, #88831, 20 µL suspension per selection condition) were prewashed with 200 µL wash buffer (20 mM HEPES, pH 7.4, 150 mM NaCl, 0.05% Tween 20, and 10 mM imidazole) for three times. The magnetic beads were then incubated with 5 µg His-tagged 3CLpro in 100 µL selection buffer (20 mM HEPES, pH 7.4, 150 mM NaCl,

0.05% Tween 20, 10 mM imidazole, 0.25 mM DTT, 0.1 mg/mL sheared salmon sperm DNA, and 0.4% DMSO) for 30 minutes at room temperature with continuous gentle rotation. A no-target control condition was performed in parallel in which no protein was added. The beads were washed with 200 μL selection buffer. The immobilized protein along with the beads were subsequently incubated with 1 nmole of pooled DNA-encoded libraries for 1 hour at room temperature in 100 μL selection buffer with continuous gentle rotation. The beads were washed with selection buffer (200 μL x3). Bound DEL compounds were recovered by heat elution in elution buffer (20 mM HEPES, pH 7.4, 150 mM NaCl, 0.05% Tween 20, and 10 mM imidazole) at 95 °C for 10 minutes.

This selection process was repeated for two rounds, with the output of the first-round selection carried forward as the input to the second-round selection with fresh protein and new beads. DEL compounds from each round were analyzed by qPCR. The coding sequences of the DEL compounds from the second-round selection were amplified by PCR and further purified by AMPure XP Beads (Beckman Coulter, #A63881). Sequencing of the PCR amplified samples was then performed on an Illumina NovaSeq 6000 instrument.

Docking and free energy calculations

Docking calculations were performed with MOE v2020 using the canonical site of 3CLpro (2) as the docking pocket. Induce fit protocol was used in which side chain atoms were allowed to move and all the single bonds in ligands were rotatable. The GBVI/WSA dG scoring function (3) was used for ranking. Free energy perturbation (FEP) calculations were performed to compute the relative binding affinities between two compounds using CHARMM/OpenMM interface (4,5). 3CLpro homodimer were used in FEP simulations with proteins described by the CHARMM36m (6) force field and ligand modeled with CGENFF (7).

3CLpro enzyme activity assay

The enzyme activity of SARS-CoV-2 3CLpro and SARS-CoV 3CLpro was evaluated using a fluorescence resonance energy transfer (FRET)-based assay. The final concentrations of 3CLpro and the substrate were 100 nM and 100 μM, respectively. In a 384-well plate (Corning, CLS3575), 10 μL of the 3CLpro (400 nM) in the reaction buffer (20 mM HEPES 7.4, 150 mM NaCl, 0.01% Triton X-100, 1 mM DTT) was mixed with 1 μL of serial dilutions of each inhibitor and incubated at room temperature for 1 hour. Then the plate was placed at 37 °C for 5 minutes, and 29 μL of the fluorogenic substrate (Dabcyl-KTSAVLQSGFRKM-Edans) in the reaction buffer (138 μM) was added to each well to initiate the reaction. The plate was immediately put into a Thermol Varioskan LUX plate reader and monitored at 37 °C using an excitation wavelength of 355 nm and an emission wavelength of 538 nm. A control experiment containing only the fluorogenic substrate in the reaction was carried out. After subtracting the control data from the experimental data, the slope of each fluorescence curve from 0 to 5 minutes was calculated as the velocity of the corresponding reaction. To calculate the $IC_{50}$ values of each inhibitor, all the velocity values were normalized to the velocity of the reaction that only DMSO was added to 3CLpro. Three independent experiments were performed. The data was analyzed using GraphPad Prism 9 software.

The enzyme activity of MERS-CoV 3CLpro was measured using a similar protocol. The differences were the final concentrations of 3CLpro and the substrate, which were 500 nM and 200 μM, respectively.

Crystallization

The 3CLpro/inhibitor complex was prepared by incubating the 3CLpro of SARS-CoV-2, SARS-CoV, or MERS-CoV (10 mg/ml in 20 mM HEPES, pH7.4, 150 mM NaCl) with 1.5 mM inhibitor at room temperature for 1 hour, followed by centrifugation at 21,000 g for 5 minutes to remove the precipitate. For crystallization, 0.2 μL of the protein/inhibitor complex was mixed with 0.2 μL of the well buffer in 96-well sitting drop crystallization plates using a protein crystallization robot (Mosquito), then the drop was equilibrated against 90 μL of the well buffer at 20 °C. The well buffer for the crystallization of SARS-CoV-2 3CLpro/WU-02 complex contained 0.05 M citric acid, 0.05 M BIS-TRIS propane pH 5.0, 16% w/v polyethylene glycol 3,350. The well buffer for the crystallization of SARS-CoV-2 3CLpro/WU-04 complex contained 1.5% v/v Tacsimate pH 4.0, 0.1 M sodium acetate trihydrate pH 4.6, 20% w/v polyethylene glycol 3,350. The well buffer for the crystallization of SARS-CoV 3CLpro/WU-04 complex contained 10% 2-propanal, 0.1 M sodium citrate pH 5.4, 10% polyethylene glycol 10,000. For the crystallization of MERS-CoV 3CLpro/WU-04 complex, the protein stock was supplemented with 10 mM DTT, and then mixed with the well buffer containing 4% v/v Tacsimate pH 5.0, 0.1M sodium citrate tribasic pH 5.4, 12% v/v polyethylene glycol 3,350, and 8 mM $CaCl_2$.

<u>Data collection and structure determination</u>

The crystals were first transferred to a cryoprotectant solution (the well buffer plus 20 mM HEPES pH 7.4, 150 mM NaCl and 20% glycerol), then loaded onto the X-ray diffractometer (Rigaku, XtaLAB Synergy Customer) at Westlake University. The diffraction data was collected at 100 K and processed with the reduction program CrysAlisPro. The structures were solved by molecular replacement using Phaser in PHENIX (8). The crystal structures of SARS-CoV-2 3CLpro (PDB code: 6Y2E), SARS-CoV 3CLpro (PDB code: 1UJ1), and MERS-CoV 3CLpro (PDB code:

4YLU) were used as the initial models. The structures were manually refined with Coot (9) and PHENIX (10). Data collection and refinement statistics can be found in table S1 that was generated using the utility phenix.table_one in PHENIX (10).

Isothermal titration calorimetry (ITC)

ITC experiments were done with the isothermal titration calorimeter MicroCal PEAQ-ITC (Malvern Panalytical). The inhibitor with a concentration of 25 μM in the ITC buffer (20 mM HEPES, pH 7.4, 150 mM NaCl, 0.625% DMSO) was titrated by 255 μM 3CLpro in the ITC buffer at 25 °C. Control experiments in which the inhibitors were titrated by the ITC buffer were performed and the data was used to correct the 3CLpro titration data. The data was processed using the MicroCal PEAQ-ITC analysis software. Each measurement was repeated three times.

Antiviral assays in multiple cell lines

***Ethics statement***

Experiments with infectious SARS-CoV-2-Nluc, SARS-CoV-2-Fluc, SARS-CoV and MERS-CoV in Vero E6 and human cells were performed at the Biosafety level-3 facility at the University of Texas Medical Branch at Galveston. Experiments with infectious SARS-CoV-2 (nCoV-SH01) in A549-hACE2 cells were conducted at the Biosafety Level 3 Laboratory of Fudan University. Experiments with infectious SARS-CoV-2 Omicron and Delta variants in Caco-2 and A549-TMPRSS2-ACE2 cells were conducted using Biosafety Level 3 facilities in Queen Mary Hospital, The University of Hong Kong.

***Cells and viruses***

African green monkey kidney epithelial Vero E6 cells (ATCC® CRL-1586), human lung adenocarcinoma epithelial Calu-3 cells (ATCC® HTB-55) , and Caco-2 cells (ATCC®HTB-37) were obtained from the American Type Culture Collection (ATCC, Bethesda, MD) and maintained in Dulbecco's modified Eagle's medium (DMEM) with 10% fetal bovine serum (FBS; HyClone Laboratories, South Logan, UT) and 1% penicillin/streptomycin (P/S; 10,000 U/mL) (Gibco). The human alveolar epithelial cell line that stably expresses human angiotensin-converting enzyme 2 (A549-hACE2) was maintained in a high-glucose DMEM supplemented with 10% fetal bovine serum, 1% P/S and 1% 4-(2-hydroxyethyl)-1-piperazineethanesulfonic acid (HEPES) and 10 μg/mL Blasticidin S (11). The primary human normal human bronchial epithelial cells were purchased from Lonza (Basel, Switzerland) and maintained in BEGM™ Bronchial Epithelial Cell Growth Medium (Lonza) at 37 °C with 5% $CO_2$ by following the manufacturer's instruction. The human ACE2 & TMPRSS2 expressing A549 cells (A549-ACE2-TMPRSS2) were purchased from Invivogen. All antibiotics and supplements were purchased from ThermoFisher Scientific (Waltham, MA). All cell lines were verified and tested negative for mycoplasma.

Two reporter viruses SARS-CoV-2-Nluc and SARS-CoV-2-Fluc used in the antiviral assays were engineered by inserting nanoluciferase (Nluc) or firefly luciferase gene at the ORF7 of the viral genome using an infectious clone of SARS-CoV-2 (strain 2019-nCoV/USA_WA1/2020) according to the protocols described previously (12,13). SARS-CoV-2 Omicron (BA.1) and Delta variants were isolated from respiratory tract specimens of laboratory-confirmed COVID-19 patients in Hong Kong (14). The sequences were deposited in GISAID with the following accession codes: Delta (hCoV-19/Hong Kong/HKU-220209-001/2021; EPI_ISL_9681416) and Omicron (hCoV-19/Hong Kong/HKU-691/2021; EPI_ISL_7138045).

***SARS-CoV-2 antiviral assays (Figs. 3a, 3b and 3c, fig. S6)***

For antiviral assay in A549-hACE2 cells, 12,000 cells per well in phenol-red free medium containing 2% FBS were plated into a white opaque 96-well plate (Corning). On the next day, 3-fold serial dilutions of compounds were prepared in dimethyl sulfoxide (DMSO). The compounds were further diluted 100-fold in the phenol-red free culture medium containing 2% FBS. Cell culture fluids were removed and incubated with 50 μL of diluted compound solutions and 50 μL of SARS-CoV-2-Nluc viruses (MOI 0.025). At 48 hours postinfection, 50 μL Nano luciferase substrates (Promega) were added to each well. Luciferase signals were measured using a Synergy™ Neo2 microplate reader. The luciferase signals (Y-axis) versus the compound concentrations were plotted in software Prism 9 and fitted using the built-in nonlinear regression method "[Inhibitor] vs. response – Variable slope (four parameters)" to get the "Bottom" and "Top" values in the following equation in which "Y" is the luciferase signal and "X" is the compound concentration:

$$Y = Bottom + (Top – Bottom)/(1+(IC50/X)^HillSlope)$$

The DMSO control was assigned a concentration of 0.1 nM. The relative luciferase signals were calculated by normalizing the luciferase signals using the following equation in which "Y" is the relative luciferase signal, "X" is the luciferase signal, "Bottom" and "Top" values are from the above equation:

$$Y = 100*(X – Bottom)/(Top – Bottom)$$

The relative luciferase signals (Y-axis) versus the compound concentrations (X-axis) were plotted in software Prism 9 and fitted using the built-in nonlinear regression method "[Agonist] vs. response – Find ECanything" to calculate the $EC_{50}$ and $EC_{90}$ values. The DMSO control was assigned a concentration of 0.1 nM during data analysis, but in the figures it was manually changed back to 0 nM.

For antiviral assay in Vero E6 and Calu-3 cells, 20,000 (Vero E6) or 30,000 (Calu-3) cells per well in medium containing 2% FBS were plated into a clear 96-well plate (Nunc). On the next day, 3-fold serial dilutions of compounds were prepared in dimethyl sulfoxide (DMSO). The compounds were further diluted 100-fold in the culture medium containing 2% FBS. Cell culture fluids were removed and incubated with 50 μL of diluted compound solutions and 50 μL of SARS-CoV-2-Fluc viruses (MOI 0.1 for Vero E6 cells, MOI 2 for Calu-3 cells). At 48 hours postinfection, cells were lyzed in 100 μL 1× cell culture lysis buffer (Promega) for 20 minutes. 40 μL lysates were mixed with 100 μL firefly luciferase substrates (Promega) in a white opaque 96-well plate (Corning), and luciferase signals were measured using a Synergy™ Neo2 microplate reader. The $EC_{50}$ and $EC_{90}$ values were calculated as above. Experiments were performed with technical duplicates.

For antiviral test in NHBE cells, $1\times10^5$ cells per well were plated into a 24-well plate (Nunc). On the next day, 3-fold serial dilutions of compounds were prepared in dimethyl sulfoxide (DMSO). The compounds were further diluted 1000-fold in the culture medium containing SARS-CoV-2-Fluc (MOI 10) and added to the cells. At 24 hours post-infection, cells were washed in PBS and lyzed in 300 μL 1× cell culture lysis buffer (Promega) for 20 minutes. 40 μL lysates were mixed with 100 μL firefly luciferase substrates (Promega) in a white opaque 96-well plate (Corning) and luciferase signals were measured using a Synergy™ Neo2 microplate reader. The $EC_{50}$ was calculated as above. Experiments were performed with six replicates.

***SARS-CoV-2 antiviral assays (Fig. 3d)***

25,000 A549-hACE2 cells per well in medium containing 10% FBS were plated into a clear 96-well plate. On the next day, 3-fold serial dilutions of compounds starting from 100 μM were prepared in the culture medium containing 2% FBS. Cell culture fluids were removed and incubated with 50 μL of diluted compound solutions and 50 μL of SARS-CoV-2 virus (nCoV-SH01, MOI 0.5). At 24 hours postinfection, cells were fixed with 4% paraformaldehyde in PBS for 30 minutes, permeablized with 0.2% Triton X-100 for 1 hour. Cells were then incubated with house-made mouse anti-SARS-CoV-2 nucleocapsid (N) protein serum (1:1000) for 2 hours at room temperature. After three washes, cells were incubated with the secondary goat anti-mouse IgG (H + L) conjugated with Alexa Fluor 555 (Thermo #A-21424, 2 μg/ml) for 1 hour at room temperature, followed by staining with 4',6-diamidino-2-phenylindole (DAPI). Images were collected using an Operetta High Content Imaging System (PerkinElmer), and processed using the PerkinElmer Harmony high-content analysis software v4.9 and ImageJ v2.0.0 (http://rsb.info.nih.gov/ij/). For each sample, the ratio of infected cells were calculated by dividing the number of N protein-positive cells by the number of nucleuses stained by DAPI; next, the inhibition of virus infection by the compounds was calculated using the following equation, in which $R_{Compound}$ and $R_{DMSO}$ were the ratios of infected cells in the presence of the compounds and DMSO, respectively:

$$\text{Inhibition (\%)} = 100*(1-R_{Compound}/R_{DMSO})$$

The Inhibition (%) versus the compound concentration (X-axis) was plotted in software Prism 9 and fitted using the built-in nonlinear regression method "[Inhibitor] vs. response – Variable slope (four parameters)" to calculate the $EC_{50}$. The DMSO control was assigned a concentration of 0.01 nM during data analysis, but in the figures it was manually changed back to 0 nM. Experiments were performed with technical duplicates.

***SARS-CoV-2 antiviral assays (Figs. 3e, 3f, 3g and 3h)***

A viral load reduction assay was performed on Caco-2 and A549-TMPRSS2-ACE2 cells, as described previously but with modifications (15). 30,000 cells/well of Caco-2 cells or A549-

TMPRSS2-ACE2 cells in phenol-red free medium containing 2% FBS were plated into a white opaque 96-well plate (Corning). On the next day, 3-fold serial dilutions of compounds were prepared in dimethyl sulfoxide (DMSO). The 3CLpro inhibitors were further diluted 100-fold in the phenol-red free culture medium containing 2% FBS. One hour after SARS-CoV-2 Omicron or Delta variant infection (MOI 0.1), the infectious inoculum were removed and replaced with 100 μL of diluted compound solutions. At 48 h.p.i., cell lysate samples from the infected cells (MOI = 0.1) were collected for qRT–PCR analysis. Briefly, 100 μl of the cell lysate were extracted for total RNA with the RNeasy Mini Kit (Qiagen). Real-time one-step qRT–PCR was used for quantitation of SARS-CoV-2 viral load, and for quantitation of the β-actin gene of the host cells that was used as an internal control, using the QuantiNova Probe RT-PCR kit (Qiagen) with a LightCycler 480 Real-Time PCR System (Roche). Each 20-μl reaction mixture contained 10 μl of 2×QuantiNova Probe RT-PCR Master Mix, 1.2 μl of RNase-free water, 0.2 μl of QuantiNova Probe RT-Mix, 1.6 μl each of 10 μM forward and reverse primer, 0.4 μl of 10 μM probe and 5 μl of extracted RNA as the template. Reactions were incubated at 45 °C for 10 min for reverse transcription, 95 °C for 5 min for denaturation, followed by 45 cycles of 95 °C for 5 s and 55 °C for 30 s. Signal detection was carried out and measurements were made in each cycle after the annealing step. The cycling profile ended with a cooling step at 40 °C for 30 s. The primers and probe for the viral gene were against the RNA-dependent RNA polymerase (RdRp) gene region of SARS-Cov-2; their sequences were as below:

forward primer: 5′ CGCATACAGTCTTRCAGGCT-3′

reverser primer: 5′ -GTGTGATGTTGAWATGACATGGTC-3′

probe: 5′ -FAM TTAAGATGTGGTGCTTGCATACGTAGAC-IABkFQ-3′

The pre-designed probe and primer set for the β-actin gene were purchased from ThermoFisher (Cat#4333762).

After qRT-PCR, the relative viral gene expression (RdRp gene copy/β-actin) in each sample was calculated using the following equation in which $Cq_{RdRp}$ and $Cq_{\beta\text{-actin}}$ were the quantitation cycles of RdRp and β-actin, respectively:

$$\text{RdRp gene copies/}\beta\text{-actin} = 2\text{^}(Cq_{RdRp} - Cq_{\beta\text{-actin}})$$

The "RdRp gene copies/β-actin" (Y-axis) versus the inhibitor concentrations were plotted in software Prism 9 and fitted using the built-in nonlinear regression method "[Inhibitor] vs. response – Variable slope (four parameters)" to get the "Top" value (the "Bottom" was set to 0) in the following equation in which "Y" is the "RdRp gene copies/β-actin" and "X" is the compound concentration:

Y = Bottom + (Top – Bottom)/(1+(IC50/X)^HillSlope)

The DMSO control was assigned a concentration of 0.01 nM. Then the "RdRp gene copies/β-actin" were normalized using the following equation:

"RdRp gene copies/β-actin (%)" = 100*"RdRp gene copies/β-actin"/Top

The "RdRp gene copies/β-actin (%)" versus the compound concentrations (X-axis) were plotted in software Prism 9 and fitted using the built-in nonlinear regression method "[Inhibitor] vs. response – Variable slope (four parameters)" to calculate the $EC_{50}$. The DMSO control was assigned a concentration of 0.01 nM during data analysis, but in the figures it was manually changed back to 0 nM.

***SARS-CoV and MERS-CoV antiviral assays***

Vero E6 cells (20,000) or Calu-3 2B4 cells (30,000) per well were seeded in a 96-well plate. The next day, WU-04 was 3-fold serially diluted in 90% dimethyl sulfoxide (DMSO). The compound dilutions were then further (x250 fold) diluted in Dulbecco's Modified Eagle Medium (DMEM) containing 2% fetal bovine serum (FBS) and mixed with an equal volume of derived SARS-CoV or MERS-CoV. The inhibitor-virus mixtures were then immediately used to infect Vero E6 cells (MOI 0.1) or Calu-3 2B4 cells (MOI 1) at 37 °C, 5% $CO_2$ for 1.5 hours. After removing the inoculum, cells were washed once with DMEM containing 2% FBS and serial dilutions of the inhibitor (500x diluted from the 90% DMSO dilutions). After an additional 24 hours (Vero E6) or 48 hours (Calu-3) of incubation at 37 °C with 5% $CO_2$, supernatants were collected and titered via plaque assay. Experiments were performed in triplicate.

Antiviral assays in mouse models

***Ethics statement***

All experiments with infectious SARS-CoV-2 were performed in the biosafety level 4 and animal biosafety level 4 facilities in the Harbin Veterinary Research Institute (HVRI) of the Chinese Academy of Agricultural Sciences (CAAS), and approved by the Committee on the Ethics of Animal Experiments of the HVRI of CAAS (approval number 211214-01).

***Cells and viruses***

Vero E6 cells were maintained in DMEM containing 10% FBS and antibiotics and incubated at 37 °C with 5% $CO_2$. Mouse-adapted SARS-CoV-2/HRB26/human/2020/CHN (HRB26M, GISAID access no. EPI_ISL_459910) and SARS-CoV-2/HRB25/human/2020/CHN (HRB25, GISAID access no. EPI_ISL_467430) were obtained as reported previously (16,17). Infectious virus titers were determined by using a standard plaque-forming unit (PFU) assay in Vero E6 cells. Viral stocks were stored in aliquots at −80 °C until use.

***Mouse studies***

Specific pathogen-free female BALB/c mice aged 6–8 weeks were obtained from Beijing Vital River Laboratory Animal Technologies Co., Ltd (Beijing, China). Male H11-K18-hACE2 transgenic mice aged 6–8 weeks were obtained from Gempharmatech Co., Ltd (Nanjing, China). Compound WU-04 and PF-07321332 were first dissolved in one volume of Tween80/ethanol (v/v = 1/1), then mixed with five volumes of 10% VE-TPGS (w/v). For BALB/c mouse study, mice (n = 6 per group) were orally administered with WU-04 (250 mg/kg of body weight per dose, twice daily) or the vehicle control for three days, and infected intranasally with 100 PFU SARS-CoV-2 HRB26M strain in a volume of 50 μL one hour after the first dose. After three days treatment, the mice were sacrificed for viral load quantification by qPCR and PFU assay in the turbinates and lungs. For H11-K18-hACE2 mice study, mice (n = 6 per group) were infected intranasally with 100 PFU SARS-CoV-2 HRB25 strain in a volume of 50 μL, and orally administered with WU-04 (100, 200, or 300 mg/kg of body weight per dose, twice daily), or PF-07321332 (300 mg/kg of body weight per dose, twice daily), or the vehicle control one hour after infection. The mice were observed daily for signs of disease, bodyweight changes for four days, and sacrificed for viral load quantification in the turbinates, lungs and brain, histopathologic analysis in the lungs, and immunohistochemical analysis in brains.

***qPCR***

Viral genomic RNA of SARS-CoV-2 was extracted by using a QIAamp vRNA Mini kit (Hilden, Germany). Reverse transcription was performed by using the HiScript II Q RT SuperMix (Nanjing, China) for qPCR. qPCR was performed to quantitate the number of viral N gene RNA copies by using the Applied Biosystems QuantStudio 5 Real-Time PCR System (Waltham, USA) with Premix Ex Taq (probe qPCR) (Dalian, China). The *N* gene-specific primers (forward, 5'-GGGGAACTTCTCCTGCTAGAAT-3'; reverse, 5'-CAGACATTTTGCTCTCAAGCTG-3') and probe (5'-FAM-TTGCTGCTGCTTGACAGATT-TAMRA-3') were utilized according to the information provided by the National Institute for Viral Disease Control and Prevention, China (http://nmdc.cn/nCoV). The amount of vRNA for the target SARS-CoV-2 *N* gene was normalized to a standard curve obtained by using a plasmid (pBluescriptIISK-N, 4,221 bp) containing the full-length cDNA of the SARS-CoV-2 *N* gene, as previously described (18).

***Histopathologic and immunohistochemical studies***

Pathology of lung was evaluated by hematoxylin and eosin (H&E) staining, and viral antigen retrieval of brain was analysis by specific anti-SARS-CoV-2 nucleoprotein monoclonal antibody staining, as previously described (19). One-µm-thick formalin-fixed paraffin-embedded sections were prepared. The slides of lung were stained and evaluated by a pathologist in a double-blind manner. The pathological scores were calculated by a pathological scoring system established by the Harbin Veterinary Research Institute (HVRI) of the Chinese Academy of Agricultural Sciences (CAAS). The slides of brain were visualized with DAB (3,3′-Diaminobenzidine) staining and counterstained with hematoxylin after immunostaining.

Preclinical pharmacokinetics studies

The preclinical pharmacokinetics studies were performed at WuXi AppTec following the study protocol and local Standard Operating Procedures (SOPs).

***Mouse pharmacokinetics***

The mice were acclimated to the test facility for at least 3 days before being placed on study. Thirty C57BL/6J mice (male, 6-9 weeks) were divided into six groups (5 mice per group), fasted overnight and given food 4 hours post dose. The mice's body weights were measured on the morning of the dosing day. Compound WU-04 and RTV were first dissolved in one volume of

Tween80/ethanol (v/v = 1/1), then mixed with five volumes of 10% VE-TPGS (w/v). A clear solution or homogenous suspension of compound WU-04 was administered via oral gavage; for the groups in which RTV was co-administered, a clear solution of RTV (2.00 mg/mL) was administered via oral gavage immediately after the administration of WU-04. Each blood sample (about 0.02 mL) was collected and processed for plasma by centrifugation. The plasma concentrations of WU-04 were determined through LC-MS/MS analysis.

***Dog pharmacokinetics***

The Beagle dogs (male, ≥ 6 months, body weight 6-12 kg) were acclimated to the test facility for at least 7 days before being placed on study. The dogs were fed twice daily (approximately 220 grams of Certified Dog Diet daily). For IV dosing, compound WU-04 was dissolved in a mixed solvent containing 5% DMSO, 5% solutol and 90% water, and then administered to four Beagle dogs via IV route. Four PO dosing, the dogs were fed the afternoon (3:30 pm to 4:00 pm) prior to the day of dosing (the remaining food was removed at night) and then fed once on the day of dosing (220 g of Certified Dog Diet, 4 hours post dose); compound WU-04 was dissolved in 10% VE-TPGS to prepare a stock of 5 mg/mL, or in a mixture of Kolliphor ELP (13.47%), Transcutol HP (66.74%) and Labrafac M1944 CS (19.79%) to prepare a stock of 200 mg/mL. For the dogs that were also administered RTV, a dose of RTV (3.5 mg/mL in 10% VE-TPGS) was administered 12 hours before the administration of WU-04, and a second dose of RTV (3.5 mg/mL in the WU-04 stock in 10% VE-TPGS) was administered together with WU-04; 12 hours later, the last dose of RTV (3.5 mg/mL in 10% VE-TPGS) was administered. Each blood collection (about 0.5 mL per time point) was performed from peripheral vein into pre-chilled commercial tube containing potassium (K2) EDTA (0.85-1.15 mg) and placed on wet ice until centrifugation. After centrifugation, the plasma concentrations of WU-04 were determined through LC-MS/MS analysis.

***Inhibition of metabolism of WU-04 in human liver microsomes by CYP isoform-selective chemical inhibitors***

The metabolism of compound WU-04 in human liver microsomes (Corning, Cat. No. 452117) was measured in the presence or absence of the CYP isoform selective inhibitors in duplicate (n = 2). To evaluate the effect of inhibition of each CYP isoform on the metabolism of WU-04, a mixture containing 100 µL of human liver microsomes (0.2 mg/mL), 10 µL of a CYP isoform selective inhibitor (potassium phosphate buffer containing 60 µM of CYP1A2 inhibitor α-Naphthoflavone, or 100 µM of CYP2B6 inhibitor Ticlopidine, or 60 µM of CYP2C8 inhibitor Montelukast, or 200 µM of CYP2C9 inhibitor Sulfaphenazole, or 40 µM of CYP2C19 inhibitor (+)-N-3-benzylnirvanol, or 40 µM of CYP2D6 inhibitor Quinidine, or 20 µM of CYP3A inhibitor Ketoconazole) or the same volume of potassium phosphate buffer, and 2 µL of compound WU-04 (100 µM) was warm up at 37 °C for 10 minutes. The reactions were initiated by addition of 88 µL of cofactor working solution (Potassium phosphate buffer containing 6.81 mM $MgCl_2$, 2.95 mM NADP, 11.4 mM glucose 6-phosphate, and 2.72 units/mL glucose-6-phosphate dehydrogenase), and after 10 minutes, the reactions were stopped by adding 600 µL of quenching solution (200ng/mL tolbutamide and 200 ng/mL labetalol in acetonitrile). As a control for each reaction, the quenching solution was added before adding the cofactor working solution. The remaining WU-04 in each reaction was analyzed by LC-MS/MS. The %Inhibition of metabolism of WU-04 was calculated using the following equation in which $R_1$ and $R_2$ represent the %Remaining of WU-04 in the presence and absence of CYP inhibitor, respectively:

$$\text{Inhibition} =.(R_1 - R_2)/(100 - R_2)$$

***Inhibition of metabolism of WU-04 in liver microsomes by ritonavir (RTV)***

The metabolism of WU-04 in human liver microsomes (Corning, Cat. No. 452117), CD-1 mouse liver microsomes (Xenotech, Cat. No. M1000), and Beagle dog liver microsomes (Xenotech, Cat. No. D1000) were measured in the presence or absence of ritonavir (RTV). To evaluate the effect of RTV on the metabolism of WU-04, a mixture containing liver microsomes (0.5 mg protein/mL), WU-04 (1 μM), RTV (1 μM) or the same volume of potassium phosphate buffer, and a NADPH regenerating system (about 1 unit/mL) was incubated at 37 °C. The reaction was stopped after 0, 5, 15, 30, 45 or 60 minutes by adding 300 μL of stop solution (cold acetonitrile containing 200 ng/mL tolbutamide and 200 ng/mL labetalol as internal standards). The remaining WU-04 in each reaction was analyzed by LC-MS/MS. The microsome clearance was calculated using the following equations:

$$C_t = C_0 * e\text{^}(-k_e * t);$$

$$T_{1/2} = 0.693/(-k_e);$$

$$CL_{int(mic)} = 0.693/T_{1/2}/\text{mg microsome protein per mL};$$

$$CL_{int(liver)} = CL_{int(mic)} * \text{mg microsomal protein/g liver weight} * \text{g liver weight/kg body weight}.$$

***Binding of WU-04 to Human, SD Rat, CD-1 Mouse, Beagle Dog, Cynomolgus Monkey Plasma***

The binding of WU-04 to plasma was measured using equilibrium dialysis. A loading matrix was prepared by diluting of a 400 μM stock of WU-04 in DMSO by the plasma matrix. To prepare the time zero ($T_0$) samples to be used for recovery determination, 50 μL of the loading matrix were transferred in triplicate to the sample collection plate, immediately matched with opposite blank buffer to obtain a final volume of 100 μL of 1:1 matrix/dialysis buffer (v/v) in each well, followed by the addition of 500 μL of stop solution (200 ng/mL of tolbutamide, 200 ng/mL of labetalol, and 50 ng/mL of metformin in acetonitrile) and then stored at 2-8°C pending further processes along with other post-dialysis samples. To load the dialysis device, 150 μL of the loading matrix and 150 μL of the dialysis buffer were transferred to the donor side and the receiver side of each well, respectively. The dialysis plate was placed in a humidified incubator at 37 °C with 5% $CO_2$ on a shaking platform that rotated slowly (about 100 rpm) for 4 hours, then 50 μL of samples were taken from both the buffer side and the matrix side, and transferred into new 96-well plates. Each sample was mixed with an equal volume of opposite blank buffer (or matrix) to reach a final volume of 100 μL, followed by the addition of 500 μL of stop solution. The mixture was vortexed and centrifuged at 4000 rpm for about 20 minutes, then 100 μL of the supernatant of each sample was removed for LC-MS/MS analysis. The single blank samples were prepared by transferring 50 μL of blank matrix to a 96 well plate and adding 50 μL of blank PBS buffer to each well. The blank plasma must match the species of plasma used in the plasma side of the well. Then the matrix-matched samples were further processed by adding 500 μL of stop solution, following the same sample processing method as the dialysis samples. The %Unbound, %Bound and %Recovery were calculated using the following equations:

$$\%Unbound = 100 \times F/T$$

$$\%Bound = 100 - \%Unbound$$

$$\%Recovery = 100 \times (F+T)/T_0$$

in which F is the analyte concentration or peak area ratio of analyte/internal standard on the buffer (receiver) side of the membrane, T is the the analyte concentration or peak area ratio of analyte/internal standard on the matrix (donor) side of the membrane, and $T_0$ is the analyte concentration or the peak area ratio of analyte/internal standard in the loading matrix sample at time zero.

### Synthesis of the inhibitors

Synthesis of compound WU-01

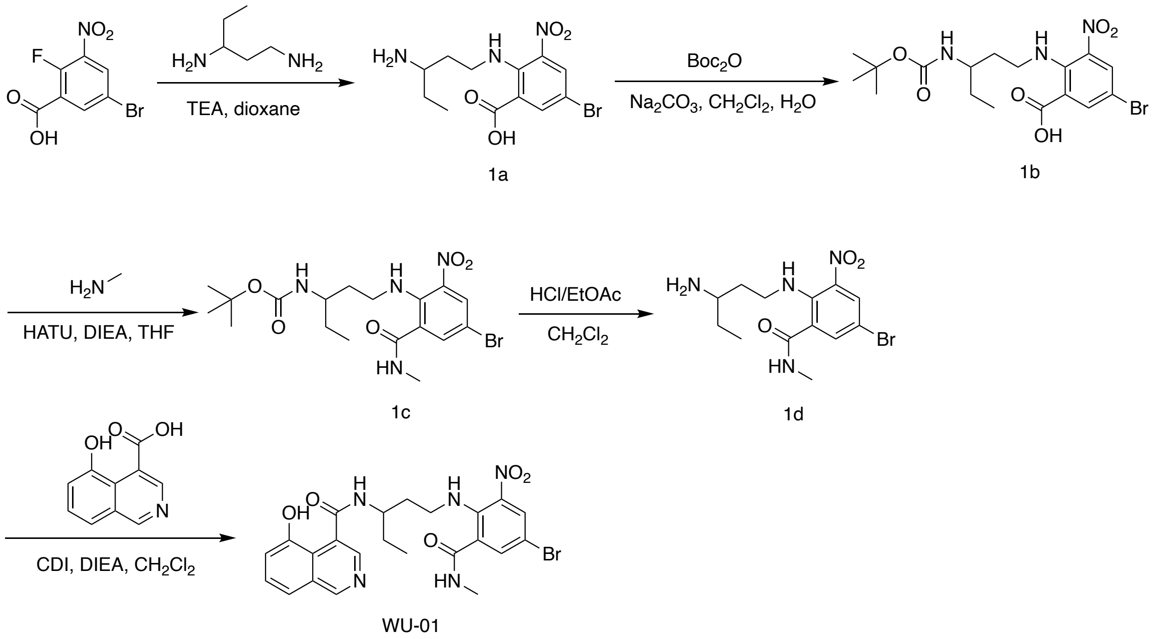


**Scheme S1. The synthesis route of compound WU-01.**

***Preparation of compound 2-(3-aminopentylamino)-5-bromo-3-nitro-benzoic acid (1a)***

To a solution of 5-bromo-2-fluoro-3-nitro-benzoic acid (500 mg, 1.89 mmol) in dioxane (5 mL), TEA (575 mg, 5.68 mmol, 790 µL) and pentane-1,3-diamine (212 mg, 2.08 mmol) were added, and the mixture was stirred at 25 °C for 1 hour. The reaction mixture was mixed with 10 mL $H_2O$ and extracted with 10 mL ethyl acetate. The organic phase was separated, washed with $H_2O$, dried over $Na_2SO_4$, and concentrated under reduced pressure to give a crude product. The crude product was purified by silica gel chromatography. Compound **1a** was obtained as a yellow solid (384 mg, 57.6% yield). LC-MS: m/z = 346.0 $[M+H]^+$.

***Preparation of compound 5-bromo-2-[3-(tert-butoxycarbonylamino) pentylamino]-3-nitro-benzoic acid (1b)***

To a solution of compound **1a** (384 mg, 1.11 mmol) in $CH_2Cl_2$ (6 mL), $H_2O$ (4 mL) and $Na_2CO_3$ (235 mg, 2.22 mmol) were added, followed by the addition of Di-tert-butyl dicarbonate ($Boc_2O$, 484 mg, 2.22 mmol) at 0 °C. Then the mixture was stirred at 20 °C for 15 hours. LC-MS showed compound **1a** was consumed completely. The reaction mixture was extracted with a mixture of 10 mL $H_2O$ and 10 mL $CH_2Cl_2$. The aqueous phase was separated, and its pH was adjusted to 6 with saturated citric acid. Then $CH_2Cl_2$ (10 mL x 3) was added to the aqueous phase to extract the product, and the organic layers were combined, dried over $Na_2SO_4$, and concentrated under reduced pressure to give a crude product. The crude product was triturated with petroleum ether at 15 °C for 15 minutes. Compound **1b** was obtained as a yellow oil (451 mg, 91.1% yield).

$^1$H NMR (400 MHz, DMSO-d6): δ 9.46-10.59 (m, 1H), 8.11 (d, J = 4.0 Hz, 1H), 7.87 (d, J = 4.0 Hz, 1H), 6.54 (d, J = 9.2 Hz, 1H), 3.09 (m, 2H), 2.69-2.81 (m, 1H), 2.56-2.65 (m, 1H), 1.32 (s, 9H), 1.18 (t, J = 7.2 Hz, 3H), 0.77 (t, J = 7.2 Hz, 3H). LC-MS: m/z = 446.0 $[M+H]^+$.

***Preparation of compound tert-butyl N-[3-[4-bromo-2-(methylcarbamoyl)-6-nitro-anilino]-1-ethyl-propyl]carbamate (1c)***

To a solution of compound **1b** (440 mg, 985.9 µmol) in THF (2 mL), methanamine hydrochloride (199 mg, 2.96 mmol), HATU (562 mg, 1.48 mmol) and DIEA (764 mg, 5.92 mmol, 1 mL) were added. The mixture was stirred at 20 °C for 15 hours. Then 10 mL $H_2O$ was added, and the mixture was extracted with 10 mL $CH_2Cl_2$. The organic phase was separated, washed with saturated $NH_4Cl$ (10 mL x 2), dried over $Na_2SO_4$, and concentrated under reduced pressure to give a crude product. The crude product was purified by silica gel chromatography. Compound **1c** was obtained as a yellow solid (97.3 mg, 21.4% yield).

$^1$H NMR (400 MHz, DMSO-d6): δ 8.62 (d, J = 4.8 Hz, 1H), 8.18 (t, J = 4.8 Hz, 1H), 8.13 (d, J = 2.4 Hz, 1H), 7.71 (d, J = 2.4 Hz, 1H), 6.56 (d, J = 8.8 Hz, 1H), 3.01-3.13 (m, 1H), 2.84-2.99

(m, 1H), 2.75 (d, J = 4.4 Hz, 3H), 1.62-1.75 (m, 1H), 1.46-1.57 (m, 1H), 1.35 (m, 1H), 1.34 (s, 9H), 1.18-1.28 (m, 2H), 0.78 (t, J = 7.2 Hz, 3H). LC-MS: m/z = 459.1 $[M+H]^+$.

***Preparation of compound 2-(3-aminopentylamino)-5-bromo-N-methyl-3-nitro-benzamide (1d)***

To a solution of compound **1c** (97.0 mg, 211 μmol) in $CH_2Cl_2$ (3 mL), 4 M HCl in ethyl acetate (4 mL) was added. The mixture was stirred at 15 °C for 4 hours, and then concentrated under reduced pressure to give compound **1d** as a yellow solid (75.0 mg, 98.9% yield).

$^1H$ NMR (400 MHz, DMSO-d6): δ 8.74-8.75 (m, 1H), 8.17 (d, J = 2.4 Hz, 1H), 8.12 (t, J = 5.2 Hz, 1H), 7.81 (br s, 3H), 7.78 (d, J = 2.4 Hz, 1H), 3.06-3.15 (m, 2H), 2.77 (d, J = 4.8 Hz, 3H), 2.31-2.37 (m, 1H), 1.74-1.87 (m, 2H), 1.47-1.58 (m, 2H), 0.88 (t, J = 7.2 Hz, 3H). LC-MS: m/z = 359.0 $[M+H]^+$.

***Preparation of compound N-[3-[4-bromo-2-(methylcarbamoyl)-6-nitro-anilino]-1-ethyl-propyl]-5-hydroxy-isoquinoline-4-carboxamide (WU-01)***

To a solution of compound 5-hydroxyisoquinoline-4-carboxylic acid (21.1 mg, 111 μmol) in DMF (4 mL), CDI (23.4 mg, 144 μmol) was added and stirred at 15 °C for 30 minutes. Then DIEA (43.1 mg, 334 μmol) and compound **1d** (40.0 mg, 111 μmol) was added and the mixture was stirred at 15 °C for 12 hours. The reaction mixture was mixed with 20 mL $H_2O$ and extracted with 40 mL $CH_2Cl_2$ (20 mL x 2). The organic layers were combined, washed with 20 mL brine, dried over $Na_2SO_4$, and concentrated under reduced pressure to give a crude product. The crude product was purified by prep-HPLC. Compound **WU-01** was obtained as a yellow solid (10.7 mg, 18.6% yield, 97.47% purity).

$^1H$ NMR (400 MHz, $CD_3OD$): δ 9.19 (s, 1H), 8.30 (s, 1H), 8.28 (d, J = 2.4 Hz, 1H), 7.66 (d, J = 2.4 Hz, 1H), 7.61-7.63 (m, 1H), 7.54-7.58 (m, 1H), 7.15-7.18 (m, 1H), 4.08-4.12 (m, 1H), 3.36-

3.25 (m, 2H), 2.89 (s, 3H), 1.95-2.01 (m, 1H), 1.77-1.80 (m, 1H), 1.61-1.67 (m, 2H), 1.05 (t, J = 7.2 Hz, 3H). LC-MS: m/z = 532.0 [M+H]$^+$.

Synthesis of compound WU-02

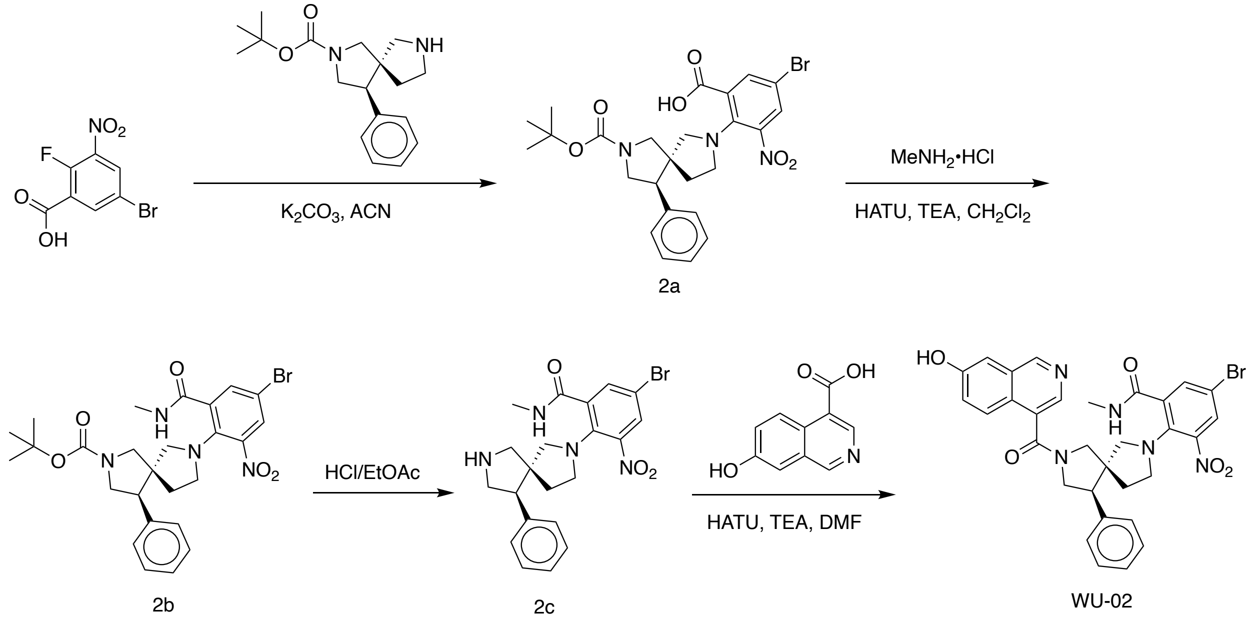


**Scheme S2. The synthesis route of compound WU-02.**

***Preparation of compound 5-bromo-2-[(4R,5S)-2-tert-butoxycarbonyl-4-phenyl-2,7-diazaspiro[4.4]nonan-7- yl] -3-nitro-benzoic acid (2a)***

To a solution of tert-butyl (4R,5S)-4-phenyl-2,7-diazaspiro[4.4]nonane-2-carboxylate (100 mg, 330 μmol) in ACN (acetonitrile, 5 mL), $K_2CO_3$ (137 mg, 992 μmol) was added while stirring at 25 °C. Then 5-bromo-2-fluoro-3-nitro-benzoic acid (175 mg, 661 μmol) was added and stirred at 25 °C for 3 hours. The mixture was filtered, and the filtrate was concentrated under vacuum to give a crude product. The crude product was purified by Prep-TLC ($CH_2Cl_2$/MeOH = 8/1) to give compound **3b** (140 mg, 75.2% yield) as a yellow solid. LC-MS: m/z = 546.1 [M+H]$^+$.

***Preparation of compound tert-butyl(4R,5S)-7-[4-bromo-2-(methylcarbamoyl)-6- nitro-phenyl]-4-phenyl-2,7-diazaspiro[4.4]nonane-2-carboxylate (2b)***

To a solution of **2a** (140 mg, 256 μmol) in $CH_2Cl_2$ (5 mL), HATU (205 mg, 512 μmol) and TEA (259 mg, 2.56 mmol, 356 μL) were added and stirred at 25 °C for 0.5 hour. Then $MeNH_2$•HCl (34.6 mg, 512 umol) was added and stirred at 25 °C for 1 hour. LC-MS showed **2a** was consumed completely. The mixture was concentrated under vacuum to give a crude product. The crude product was purified by Prep-TLC to give compound **2b** as a yellow solid (120 mg, 78.7% yield). LC-MS: m/z = 559.0 $[M+H]^+$.

***Preparation of compound 5-bromo-N-methyl-3-nitro-2-[(5R,9R)-9-phenyl-2,7-diazaspiro [4.4]nonan-2-yl]benzamide (2c)***

Compound **2b** (120 mg, 214 μmol) was dissolved in HCl/ethyl acetate (4 M, 5 mL) and stirred at 30 °C for 2 hours. LC-MS showed **2b** was consumed completely. The mixture was concentrated under vacuum to give a crude product of compound **2c** as a yellow solid (90 mg, 88.6% yield). This crude product was used in next step reaction without further purification. LC-MS: m/z = 460.9 $[M+H]^+$.

***Preparation of compound 5-bromo-2-((5S,9R)-7-(7-hydroxyisoquinoline-4-carbonyl)-9-phenyl-2,7-diazaspiro[4.4]nonan-2-yl)-N-methyl-3-nitrobenzamide (WU-02)***

To a solution of 7-hydroxyisoquinoline-4-carboxylic acid (55.6 mg, 293 μmol) in $CH_2Cl_2$ (5 mL), HATU (223 mg, 587 μmol) and TEA (297 mg, 2.94 mmol, 409 μL) were added and stirred at 30 °C for 0.5 hour. Then compound **2c** (90 mg, 195 μmol) was added and the reaction was stirred at 30 °C for 2 hours. The mixture was concentrated under vacuum to give a crude product. The crude product was purified by reversed-phase HPLC to give compound **WU-02** as a yellow solid (13.0 mg, 10.2% yield, 97% purity).

$^1$H NMR (400 MHz, $CD_3OD$): δ 9.18-9.37 (m, 1H), 8.42 (d, J = 3.2 Hz, 1H), 7.88-8.10 (m, 2H), 7.65-7.83 (m, 1H), 7.53-7.64 (m, 1H), 7.38-7.51 (m, 2H), 7.30-7.36 (m, 2H), 7.18 (d, J = 7.2 Hz, 1H), 5.82-6.50 (m, 1H), 4.06-4.28 (m, 1H), 3.87-3.95 (m, 1H), 3.46-3.74 (m, 2H), 3.34-3.42 (m, 2H), 2.93-3.22 (m, 3H), 2.87-2.91 (m, 1H), 2.47-2.68 (m, 2H), 1.42-1.80 (m, 2H). LC-MS: m/z = 632.3 $[M+2]^+$.

Synthesis of compound WU-03

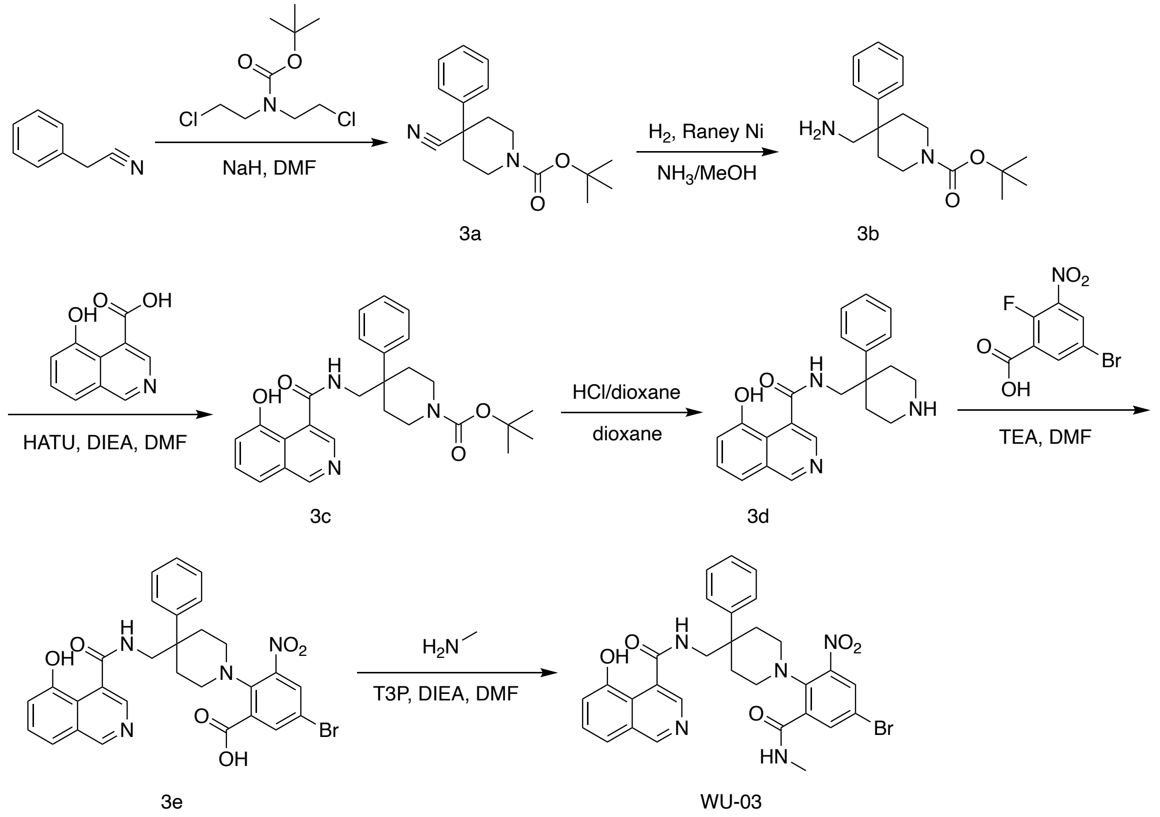


**Scheme S3. The synthesis route of compound WU-03.**

***Preparation of compound tert-butyl 4-cyano-4-phenyl-piperidine-1-carboxylate (3a)***

To a solution of 2-phenylacetonitrile (3.0 g, 25.6 mmol, 2.97 mL) and tert-butyl N,N-bis(2-chloroethyl)carbamate (6.20 g, 25.6 mmol) in DMF (40 mL), NaH (5.12 g, 128 mmol, 60% purity) was added portion wisely at 0 °C over 1 hour. The reaction mixture was stirred at 60 °C for 12 hrs. The reaction mixture was poured into ice-water (500 mL) and extracted with ethyl acetate (200 mL x 3). The organic layers were combined, washed with brine (200 mL x 2), dried over $Na_2SO_4$, filtered. After removing the solvent by vacuum evaporation, the residue was purified by flash silica gel chromatography to give compound **3a** as a yellow solid (5.0 g, 64.9% yield).

$^1$H NMR: EB2017-49-P1N2 (400 MHz, CDCl3): δ 7.30-7.56 (m, 5H), 4.29 (br d, J = 2.0 Hz, 2H), 3.22 (br s, 2H), 2.05-2.26 (m, 2H), 1.89-2.02 (m, 2H), 1.49 (s, 9H). LC-MS: m/z = 187.0 $[M-100+H]^+$.

***Preparation of compound tert-butyl 4-(aminomethyl)-4-phenyl-piperidine-1-carboxylate (3b)***

Compound **3a** (5.0 g, 17.4 mmol) was mixed with a solution of ammonia in methanol (7 M, 50 mL). After addition of Raney-Ni (2.3 g) under $N_2$, the suspension was degassed under vacuum and purged with $H_2$ several times. The mixture was stirred under $H_2$ (15 psi) at 20 °C for 12 hours, then filtered to remove the Raney-Ni. The filtrate was concentrated under vacuum to give compound **3b** as a green oil (4.7 g, crude). LC-MS: m/z = 235.2 $[M-56+H]^+$.

***Preparation of compound tert-butyl 4-[[(5-hydroxyisoquinoline-4-carbonyl)amino]methyl]-4-phenyl-piperidine-1-carboxylate (3c)***

To a solution of 5-hydroxyisoquinoline-4-carboxylic acid (50.0 mg, 264 µmol) in DMF (3 mL), HATU (120 mg, 317 µmol) and DIEA (102 mg, 792 µmol, 138 µL) was added, followed by the addition of a solution of compound **3b** (92.1 mg, 317 µmol) in DMF. The mixture was stirred at 20 °C for 1 hour. LC-MS showed that 5-hydroxyisoquinoline-4-carboxylic acid was consumed completely. The product was precipitated by adding $H_2O$ to the reaction mixture and collected by filtration under reduced pressure. Compound **3c** was obtained as a gray solid (75.0 mg, crude). LC-MS: m/z = 462.2 $[M+H]^+$.

***Preparation of compound 5-hydroxy-N-[(4-phenyl-4-piperidyl)methyl]isoquinoline-4-carboxamide (3d)***

To a solution of compound **3c** (75 mg, 162 μmol) in dioxane (2 mL), HCl/dioxane (4 M, 1.5 mL) was added. The mixture was stirred at 20 °C for 30 min, and then evaporated under reduced pressure to a crude product of compound 3d as a yellow solid (100 mg, HCl salt). LC-MS: m/z = 362.2 $[M+H]^+$.

***Preparation of compound 5-bromo-2-[4-[[(5-hydroxyisoquinoline-4-carbonyl)amino]methyl]-4-phenyl-1-piperidyl]-3-nitro-benzoic acid (3e)***

To a solution of 5-bromo-2-fluoro-3-nitro-benzoic acid (73.0 mg, 276 μmol), compound **3d** (100 mg, 276 μmol) in DMF (2 mL) and TEA (83.9 mg, 830 μmol, 115 μL) were added. The mixture was stirred at 20 °C for 12 hours. LC-MS showed that 5-bromo-2-fluoro-3-nitro-benzoic acid was consumed completely. The reaction mixture was mixed with $CH_2Cl_2$ (10 mL) and washed with $H_2O$ (5 mL x 3). The organic layer was dried over $Na_2SO_4$ and filtered. After removing the solvent under reduced pressure, the residue was purified by prep-HPLC to give compound **3e** as a yellow solid (50.0 mg, 29.8% yield). LC-MS: m/z = 607.1 $[M+H]^+$.

***Preparation of compound N-[[1-[4-bromo-2-(methylcarbamoyl)-6-nitro-phenyl]-4-phenyl-4-piperidyl]methyl]-5-hydroxy-isoquinoline-4-carboxamide (WU-03)***

To a solution of compound **3e** (50.0 mg, 82.5 μmol) and methanamine hydrochloride (16.7 mg, 247 μmol) in $CH_2Cl_2$ (2 mL), T3P (78.8 mg, 123 μmol, 73.6 μL, 50% purity) and DIEA (74.7 mg, 578 μmol, 100 μL) were added at 0 °C. The mixture was stirred at 20 °C for 12 hours. LC-MS showed that compound **3e** was consumed completely. After removing the solvent under reduced pressure, the residue was purified by prep-HPLC to give compound **WU-03** as a yellow solid (12.7 mg, 24.6% yield, 98.9% purity, formic acid salt).

$^1$H NMR (400 MHz, $CD_3OD$): δ 9.34 (br s, 1H), 8.54 (br s, 1H), 8.39 (d, J = 2.4 Hz, 1H), 8.31 (s, 1H), 8.18 (d, J = 2.43 Hz, 1H), 7.92-7.94 (m, 1H), 7.59 (t, J = 8.0 Hz, 1H), 7.36-7.51 (m, 5H), 7.30 (br t, J = 6.8 Hz, 1H), 7.02 (d, J = 7.6 Hz, 1H), 3.75 (br d, J = 13.2 Hz, 1H), 3.34-3.53 (m, 3H), 2.89-3.02 (m, 2H), 2.44-2.55 (m, 2H), 2.22 (s, 3H), 2.06-2.18 (m, 2H). LC-MS: m/z = 618.4 $[M+H]^+$.

Synthesis of compound WU-04

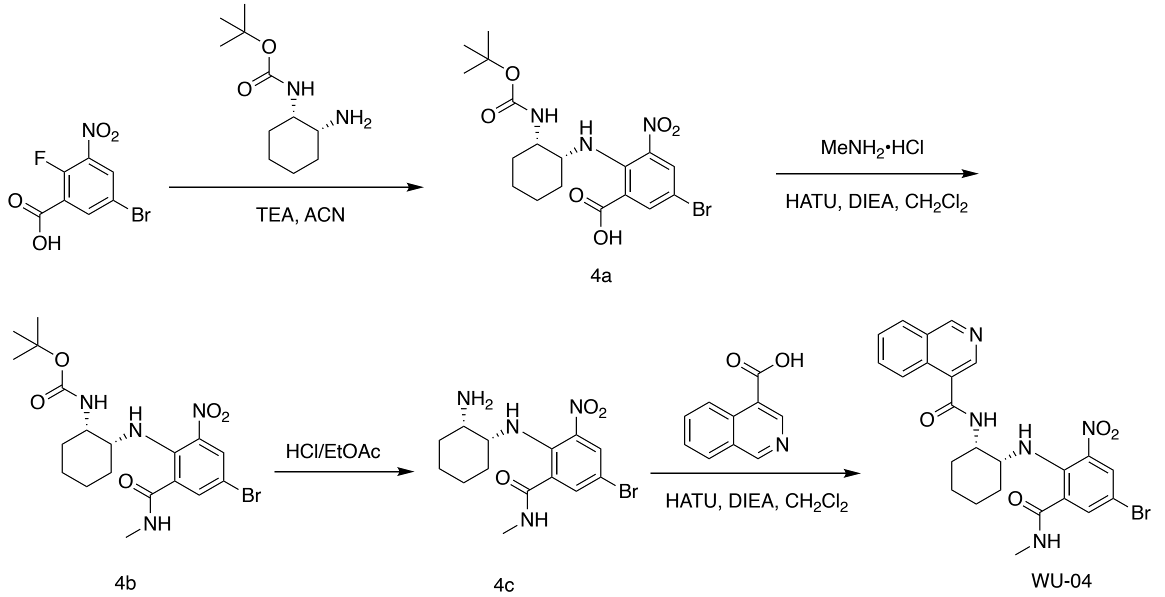


**Scheme S4. The synthesis route of compound WU-04.**

***Preparation of compound 5-bromo-2-[[(1R, 2S)-2-(tert-butoxycarbonylamino) cyclohexyl] amino]-3-nitro-benzoic acid (4a)***

To a solution of 5-bromo-2-fluoro-3-nitro-benzoic acid (200 mg, 758 μmol) in ACN (3 mL), TEA (153 mg, 1.52 mmol, 211 μL) was added, followed by the addition of tert-butyl N-[(1S, 2R)-2-aminocyclohexyl] carbamate (81.2 mg, 379 μmol). The mixture was stirred at 25 °C for 2 hours. The reaction mixture was filtered through a gelite pad, and the filtrate was concentrated to remove the solvent. The crude product was purified by Perp-TLC to give compound **4a** as a yellow solid (90.0 mg, 50.3% yield).

$^{1}$H NMR (400MHz, $CDCl_3$): δ 8.16-8.47 (m, 1H), 7.75-8.01 (m, 1H), 3.60-3.71 (m, 1H), 3.25-3.37 (m, 1H), 1.55-1.69 (m, 4H), 1.40-1.53 (m, 4H), 1.28-1.34 (m, 9H). LC-MS: m/z = 459.9 $[M+H]^+$.

***Preparation of compound tert-butyl ((1S, 2R)-2-((4-bromo-2-(methylcarbamoyl)-6-nitrophenyl) amino) cyclohexyl) carbamate (4b)***

To a solution of compound **4a** (90.0 mg, 196 umol) in $CH_2Cl_2$ (3 mL), HATU (112 mg, 295 μmol) and DIEA (140 mg, 1.08 mmol, 188 μL) were added, followed by the addition of methanamine hydrochloride (33.2 mg, 491 μmol). Then the reaction was stirred at 25 °C for 12 hours. LC-MS showed that **4a** was consumed completely. The reaction mixture was filtered through a gelite pad, and the filtrate was concentrated to remove the solvent. The crude product was purified by Perp-TLC. Compound **4b** was obtained as a yellow solid (63.6 mg, 66.0% yield).

$^{1}$H NMR (400 MHz, $CDCl_3$): δ 8.27-8.31 (m, 1H), 7.77 (d, J =2.4 Hz, 1H), 7.52-7.59 (m, 1H), 6.76-6.85 (m, 1H), 4.56-4.57 (m, 1H), 3.83 (br s, 1H), 3.65 (br s, 1H), 3.00 (d, J =4.8 Hz, 3H), 1.58-1.61 (m, 2H), 1.50-1.57 (m, 2H), 1.42-1.50 (m, 2H), 1.38-1.42 (m, 1H), 1.37 (s, 9H), 1.20-1.33 (m, 1H). LC-MS: m/z = 471.2 $[M+H)^+$.

***Preparation of compound 2-(((1R, 2S)-2-aminocyclohexyl) amino)-5-bromo-N-methyl-3-nitrobenzamide (4c)***

Compound **4b** (63.6 mg, 135 μmol) in HCl/ethyl acetate (4 M, 3 mL) was stirred at 25 °C for 15 min. LC-MS showed the **4d** was consumed completely. The reaction mixture was filtered through a gelite pad. The filtrate was concentrated to obtain compound **4c** as a yellow solid (50.0 mg, 97.8% yield). LC-MS: m/z = 370.9 $[M+H]^+$.

***Preparation of compound N-((1S,2R)-2-((4-bromo-2-(methylcarbamoyl)-6-nitrophenyl)amino)cyclohexyl)isoquinoline-4-carboxamide (WU-04)***

To a mixture of isoquinoline-4-carboxylic acid (11.7 mg, 67.3 μmol) in $CH_2Cl_2$ (3 mL), HATU (38.4 mg, 101 μmol) and DIEA (52.2 mg, 404 μmol, 70.4 μL) were added, followed by the addition of compound **4c** (50.0 mg, 135 μmol). The reaction was stirred at 25 °C for 12 hours. LC-MS showed compound **4c** was consumed completely. The reaction mixture was filtered through a gelite pad. The filtrate was concentrated to remove the solvent. The crude product was purified by Prep-HPLC to give compound **WU-04** as a yellow solid (11.0 mg, 30.5% yield, 98.75% purity).

$^1$H NMR (400 MHz, MeOD): δ (ppm) 9.38-9.44 (m, 1H), 8.65-8.79 (m, 1H), 8.20-8.33 (m, 2H), 8.16 (d, J =8.4 Hz, 1H), 7.90-7.99 (m, 1H), 7.79-7.85 (m, 1H), 7.68 (d, J =2.4 Hz, 1H), 4.41-4.45 (m, 1H), 3.97 (br d, J =4.0 Hz, 1H), 2.91-2.95 (m, 3H), 1.84-1.92 (m, 1H), 1.75-1.83 (m, 3H), 1.61-1.73 (m, 2H), 1.49-1.59 (m, 2H). LC-MS: m/z = 528.2 $[M+H]^+$.

<u>Synthesis of compound WU-05</u>

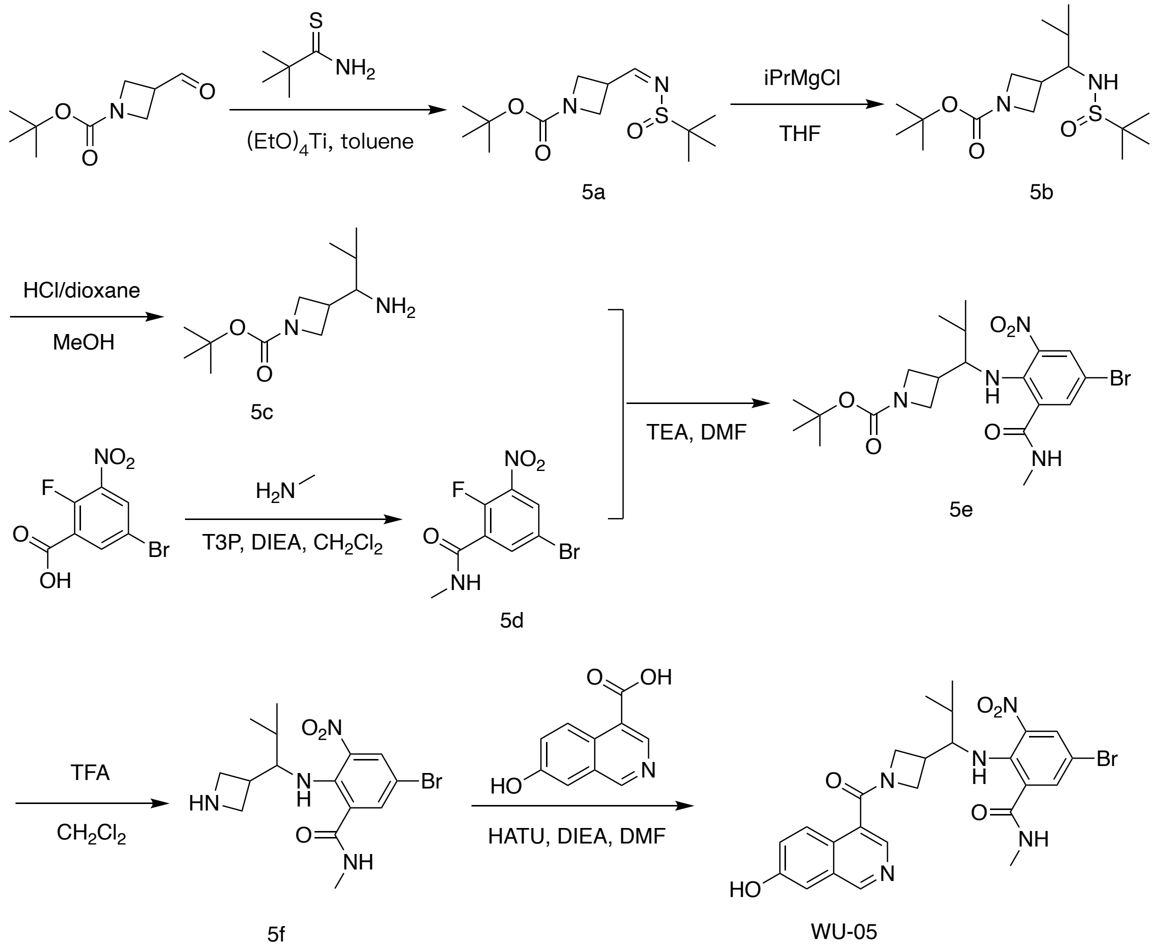


**Scheme S5. The synthesis route of compound WU-05.**

***Preparation of compound tert-butyl 3-[(Z)-tert-butylsulfinyliminomethyl]azetidine-1-carboxylate (5a)***

A mixture of 2-methylpropane-2-sulfinamide (2.16 g, 17.8 mmol), tert-butyl 3-formylazetidine-1-carboxylate (3.0 g, 16.2 mmol), and tetraethoxytitanium (7.39 g, 32.3 mmol, 6.72 mL) in toluene (6 mL) was stirred at 120 °C for 12 hours under $N_2$ protection. After removing the solvent under reduced pressure, the residue was diluted with 50 mL ethyl acetate and 20 mL $H_2O$ and filtered. The filtrate was extracted with ethyl acetate (10 mL x 3). The organic layers were combined, dried over $Na_2SO_4$, and filtered. The solvent was removed under reduced pressure and the residue was purified by flash silica gel chromatography to give compound **5a** as a yellow oil (3.0 g, 64.22% yield). LC-MS: m/z = 233.1 $[M-56+H]^+$.

***Preparation of compound tert-butyl 3-[1-(tert-butylsulfinylamino)-2-methyl-propyl]azetidine-1-carboxylate (5b)***

To a solution of compound **5a** (3.0 g, 10.4 mmol) in THF (15 mL), chloro(isopropyl)magnesium (2 M, 6.24 mL) was added dropwise at 0 °C over 10 minutes. The mixture was stirred at 20 °C for 4 hours, then poured into saturated aqueous $NH_4Cl$ (100 mL) and stirred for 20 minutes. The mixture was extracted with $CH_2Cl_2$ (10 mL x 3). The organic layers were combined, washed with saturated aqueous NaCl (10 mL x 2), dried with anhydrous $Na_2SO_4$, and filtered. After removing the solvent under reduced pressure, the residue was purified by prep-HPLC to give compound **5b** as a yellow oil (200 mg, 5.78% yield). LC-MS: m/z = 333.0 $[M+H]^+$.

***Preparation of compound tert-butyl 3-(1-amino-2-methyl-propyl)azetidine-1-carboxylate (5c)***

To a solution of compound 5b (200 mg, 601 µmol) in MeOH (20 mL), HCl/dioxane (2 M, 2 mL) was added. The mixture was stirred at 0 °C for 1 hour, then the pH was adjusted to 7 using 1 M NaOH. The mixture was filtered, and the filtrate was concentrated under reduced pressure to give compound **5c** as a yellow oil (150 mg, crude, HCl salt). LC-MS: m/z = 173.2 $[M-56+H]^+$.

***Preparation of compound 5-bromo-2-fluoro-N-methyl-3-nitro-benzamide (5d)***

To a solution of 5-bromo-2-fluoro-3-nitro-benzoic acid (500 mg, 1.89 mmol) in $CH_2Cl_2$ (6 mL), DIEA (856 mg, 6.63 mmol, 1.15 mL) and T3P (2.41 g, 3.79 mmol, 2.25 mL, 50% purity) were added, followed by the addition of a solution of methanamine hydrochloride in $CH_2Cl_2$ (191 mg, 2.84 mmol). The mixture was stirred at 20 °C for 1 hour. LC-MS showed that 5-bromo-2-fluoro-3-nitro-benzoic acid was consumed completely. Then $H_2O$ was added to the mixture to precipitate the product. Compound **5d** was obtained as a yellow solid (485 mg, 92.43% yield). LC-MS: m/z = 276.7 $[M+H]^+$.

***Preparation of compound tert-butyl 3-[1-[4-bromo-2-(methylcarbamoyl)-6-nitro-anilino]-2-methyl-propyl]azetidine-1-carboxylate (5e)***

To a solution of compound **5c** (150 mg, 656 μmol) and compound **5d** (182 mg, 656 μmol) in DMF (3 mL), TEA (199 mg, 1.97 mmol, 274 μL) was added. The mixture was stirred at 25 °C for 12 hours. Then the solvent was removed under vacuum to give a crude product. The crude product was purified by prep-HPLC. Compound **5e** was obtained as a red oil (150 mg, 44.4% yield). LC-MS: m/z = 430.9 $(M-56+3H)^+$.

***Preparation of compound 2-[[1-(azetidin-3-yl)-2-methyl-propyl]amino]-5-bromo-N-methyl-3-nitro-benzamide (5f)***

To a solution of compound **5e** (150 mg, 309 μmol) in $CH_2Cl_2$ (3 mL), TFA (1.54 g, 13.5 mmol, 1 mL) was added, and the mixture was stirred at 20 °C for 1 hour. LC-MS showed that compound **5e** was consumed completely. After removing the solvent under vacuum, a crude product of compound **5f** was obtained as a yellow oil (160 mg, crude, TFA salt). LC-MS: m/z = 386.9 $[M+H]^+$.

***Preparation of compound 5-bromo-2-[[1-[1-(7-hydroxyisoquinoline-4-carbonyl)azetidin-3-yl]-2-methyl-propyl]amino]-N-methyl-3-nitro-benzamide (WU-05)***

To a solution of 7-hydroxyisoquinoline-4-carboxylic acid (70.0 mg, 370 μmol) in DMF (3 mL), HATU (168 mg, 444 μmol) and DIEA (239 mg, 1.85 mmol, 322 μL) were added, followed by the addition of compound **5f** (156 mg, 407 μmol) in DMF (2 mL). After stirring at 20 °C for 30 minutes, the solvent was removed under vacuum and the residue was purified by prep-HPLC to give compound **WU-05** as a yellow solid (13.4 mg, 6.48% yield, 99.5% purity).

$^1$H NMR (400 MHz, $CD_3OD$): δ 9.08 (br d, J = 6.4Hz, 1H), 8.15-8.43 (m, 2H), 7.90-8.04 (m, 1H), 7.62 (dd, J = 8.8, 2.4 Hz, 1H), 7.38-7.51 (m, 1H), 7.35 (dd, J = 6.0, 2.4 Hz, 1H), 4.20-4.49 (m, 2H), 4.00-4.18 (m, 2H), 3.60-3.80 (m, 1H), 3.00-3.16 (m, 1H), 2.90 (d, J = 14.0 Hz, 3H), 1.70-1.90 (m, 1H), 0.76-0.90 (m, 6H). LC-MS: m/z = 558.3 $[M+H]^+$.

Synthesis of compound WU-06

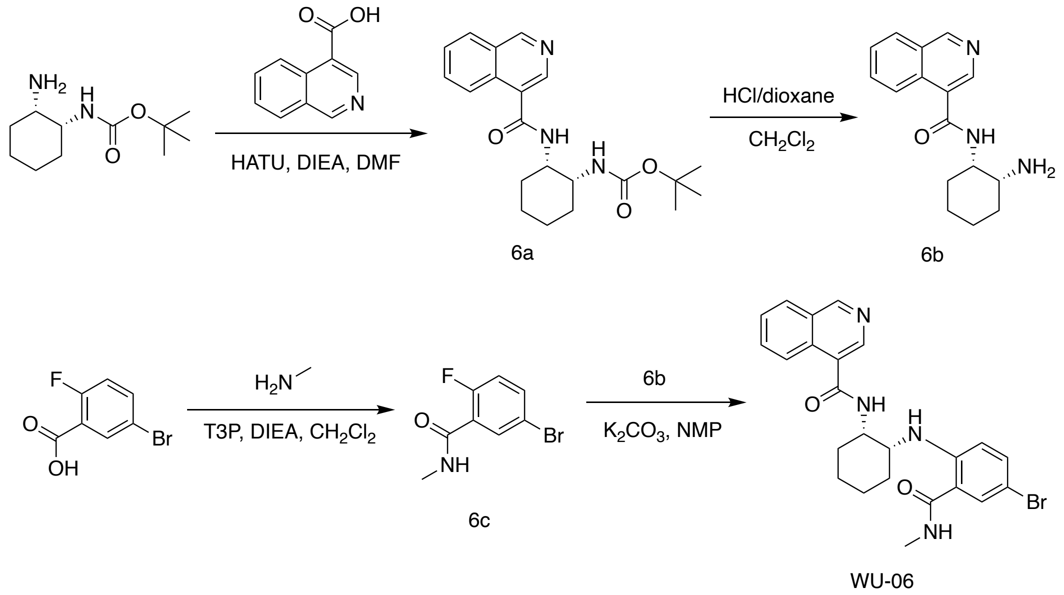


**Scheme S6. The synthesis route of compound WU-06.**

***Preparation of compound tert-butyl N-[(1R,2S)-2-(isoquinoline-4-carbonylamino) cyclohexyl]carbamate (6a)***

To a solution of isoquinoline-4-carboxylic acid (2.0 g, 11.6 mmol) in DMF (40 mL), HATU (8.78 g, 23.1 mmol) and DIEA (2.99 g, 23.1 mmol, 4 mL) were added. The mixture was stirred at 25 °C for 0.5 hour. Then tert-butyl N-[(1R,2S)-2-aminocyclohexyl]carbamate (2.97 g, 13.9 mmol) was added. The mixture was stirred 25 °C for additional 2 hours, and then poured into 50 mL water and extracted with ethyl acetate (30 mL x 2). The organic layers were combined, washed with saturated aqueous NaCl (100 mL x 2), dried with anhydrous $Na_2SO_4$. The solvent was removed under vacuum, and the residue was purified by flash silica gel chromatography. Compound **6a** was obtained as a white solid (3.6 g, 77.9% yield).

$^1$H NMR (400 MHz, $CD_3OD$): δ 9.30 (s, 1H), 8.57 (s, 1H), 8.28 (d, J = 8.0 Hz, 1H), 8.16 (d, J = 8.0 Hz, 1H), 7.83-7.90 (m, 1H), 7.71-7.78 (m, 1H), 4.27-4.38 (m, 1H), 4.06-4.18 (m, 1H), 1.62-1.80 (m, 6H), 1.49-1.55 (m, 2H), 1.40 (s, 9H). LC-MS: m/z = 370.3 $[M+H]^+$.

***Preparation of compound N-[(1S,2R)-2-aminocyclohexyl]isoquinoline-4-carboxamide (6b)***

To a solution of compound **6a** (400 mg, 1.08 mmol) in $CH_2Cl_2$ (6 mL), HCl/dioxane (4 M, 4 mL) was added. The mixture was stirred at 25 °C for 0.6 hour, and the pH was adjusted to above 7 by NaOH (3 M in $H_2O$). Then ethyl acetate (10 mL x 3) was added into the mixture to extract the product. The ethyl acetate layers were combined, washed with saturated aqueous NaCl (50 mL x 2) and dried over $Na_2SO_4$. The solvent was removed under reduced pressure to give compound **6b** as a white solid (170 mg, 55.9% yield).

$^1$H NMR: (400 MHz, $CD_3OD$): δ 9.32 (s, 1H), 8.64 (s, 1H), 8.25 (d, J = 8.0 Hz, 1H), 8.18 d, J = 8.0 Hz, 1H), 7.84-7.93 (m, 1H), 7.72-7.80 (m, 1H), 4.26-4.37 (m, 1H), 3.15-3.23 (m, 1H), 1.48-1.89 (m, 8H). LC-MS: m/z = 270.1 $[M+H]^+$.

***Preparation of compound 5-bromo-2-fluoro-N-methyl-benzamide (6c)***

To a solution of 5-bromo-2-fluoro-benzoic acid (1 g, 4.57 mmol) in DMF (5 mL), methanamine (462 mg, 6.85 mmol), TEA (1.85 g, 18.2 mmol, 2.54 mL) and T3P (5.81 g, 9.13 mmol, 5.43 mL, 50% purity) were added. The mixture was stirred at 25 °C for 4 hours, then HCI (1N) was used to adjust the pH to 7. The mixture was extracted with ethyl acetate (20 mL x 2), and the organic layers were combined, washed with saturated aqueous NaCl (50 mL x 3), dried with anhydrous $Na_2SO_4$, and filtered. The solvent was removed under reduced pressure to give compound **6c** as yellow solid (950 mg, 37.1% yield). LC-MS: m/z = 232.0 $[M+H]^+$.

***Preparation of compound 4 N-[(1S,2R)-2-[4-bromo-2-(methylcarbamoyl)anilino]cyclohexyl] isoquinoline-4-carboxamide (WU-06)***

To a solution of compound **6b** (295 mg, 1.10 mmol) and compound **6c** (170 mg, 732 μmol) in NMP (4 mL), $K_2CO_3$ (202 mg, 1.46 mmol) was added. The mixture was stirred at 120 °C for 24 hours. Then 10 mL $H_2O$ was added, and the mixture was extracted with $CH_2Cl_2$ (10 mL), washed with saturated aqueous NaCl (10 mL x 2), dried over $Na_2SO_4$. The solvent was removed under reduced pressure and the residue was purified by Prep-HPLC to give compound **WU-06** as a yellow solid (21.9 mg, 5.99% yield, 96.47% purity).

$^1$H NMR (400 MHz, $CD_3OD$): δ 9.25 (s, 1H), 8.37 (s, 1H), 8.09-8.16 (m, 2H), 7.69-7.78 (m, 2H), 7.56-7.64 (m, 1H), 7.24-7.37 (m, 1H), 6.82-6.92 (m, 1H), 4.40-4.48 (m, 1H), 4.18-4.28 (m, 1H), 2.76-2.85 (m, 3H), 1.88 (m, 5H), 1.54-1.66 (m, 3H). LC-MS: m/z = 481.0 $[M+H]^+$.

Synthesis of compound WU-07

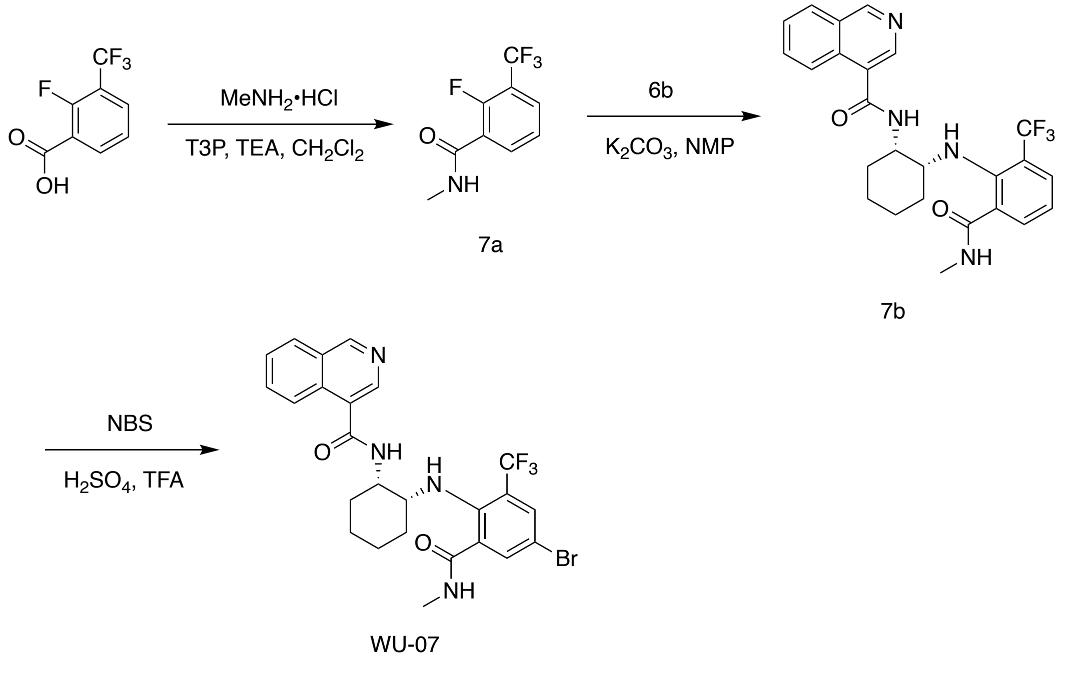

**Scheme S7. The synthesis route of compound WU-07.**

***Preparation of compound 2-fluoro-N-methyl-3-(trifluoromethyl)benzamide (7a)***

To a solution of 2-fluoro-3-(trifluoromethyl) benzoic acid (3.00 g, 14.4 mmol) in $CH_2Cl_2$ (40 mL), TEA (4.37 g, 43.2 mmol, 6.01 mL) and methanamine hydrochloride (2.43 g, 36.0 mmol) were added, followed by the addition of T3P (18.3 g, 28.8 mmol, 17.1 mL, 50% purity) at 0 °C. The mixture was stirred at 25 °C for 2 hours. LC-MS showed that the 2-fluoro-3-(trifluoromethyl) benzoic acid was consumed completely. The reaction mixture was diluted with $CH_2Cl_2$ (20 mL) and washed with $H_2O$ (50 mL x 3). The organic layers were combined and further washed with saturated aqueous NaCl (50 mL x 3), dried over $Na_2SO_4$, and filtered. The solvent was removed by evaporation under reduced pressure, and the residue was purified by flash silica gel chromatography to give compound **7a** as a yellow solid (3 g, 91.3% yield).

$^1$H NMR (400 MHz, $CDCl_3$): δ 8.24-8.37 (m, 1H), 7.74 (t, J = 6.8 Hz, 1H), 7.37 (t, J = 7.6 Hz, 1H), 6.68 (s, 1H), 3.06 (dd, J = 4.8, 0.8 Hz, 3H). LC-MS: m/z = 221.9 $[M+H]^+$.

***Preparation of compound N-[(1S, 2R)-2-[2-(methylcarbamoyl)-6-(trifluoromethyl) anilino]cyclohexyl]isoquinoline-4-carboxamide (7b)***

To a solution of compound **7a** (500 mg, 2.26 mmol) and N-[(1S,2R)-2-aminocyclohexyl]isoquinoline-4-carboxamide (compound **6b**, 608 mg, 2.26 mmol) in NMP (10 mL), $K_2CO_3$ (781 mg, 5.65 mmol) was added, and the mixture was stirred at 120 °C for 12 hours. The reaction mixture was diluted with ethyl acetate (100 mL) and washed with $H_2O$ (150 mL x 3). The organic layers were combined, washed with saturated aqueous NaCl (150 mL x 3), dried over $Na_2SO_4$, and filtered. The solvent was removed by evaporation under reduced pressure, and the residue was purified by flash silica gel chromatography to give compound **7b** as a brown solid (65 mg, 3.17% yield). LC-MS: m/z = 471.2 $[M+H]^+$.

***Preparation of compound N-[(1S, 2R)-2-[4-bromo-2-(methylcarbamoyl)-6-(trifluoromethyl) anilino] cyclohexyl] isoquinoline-4-carboxamide (WU-07)***

To a solution of compound **7b** (65 mg, 138 µmol) in a mixture of $H_2SO_4$ (0.6 mL) and TFA (0.4 mL), NBS (29.5 mg, 165 µmol) was added. The reaction was stirred at 40 °C for 1 hour. LC-MS showed that compound **7b** was consumed completely. Then the reaction mixture was added by dropwise into $H_2O$ (10 mL), followed by the addition of saturated aqueous $NaHCO_3$ to adjust the pH to 6-7. After adding 20 mL $CH_2Cl_2$, the mixture was washed with $H_2O$ (25 mL x3). The organic layers were combined, washed with saturated NaCl solution (25 mL x 3), dried over $Na_2SO4$, and filtered. The solvent was removed by evaporation under reduced pressure, and the residue was purified by prep-HPLC to give compound **WU-07** as a yellow solid (10 mg, 12.9% yield, 98.08% purity).

$^1$H NMR (400 MHz, MeOD): δ 9.32 (s, 1H), 8.54 (s, 1H), 8.16 (t, J = 7.2 Hz, 2H), 7.85 (td, J = 7.6, 1.2 Hz, 1H), 7.76 (t, J = 7.2 Hz, 1H), 7.71 (s, 1H), 7.64 (s, 1H), 4.47-4.54 (m, 1H), 3.84 (s, 1H), 2.88 (s, 3H), 1.90-1.97 (m, 1H), 1.76 (dd, J = 8.8, 4.2 Hz, 1H), 1.66-1.72 (m, 3H), 1.52-1.58 (m, 1H), 1.38-1.46 (m, 1H), 1.31 (d, J = 18 Hz, 1H). LC-MS: m/z = 549.3 $[M+H]^+$.

Synthesis of compound WU-08

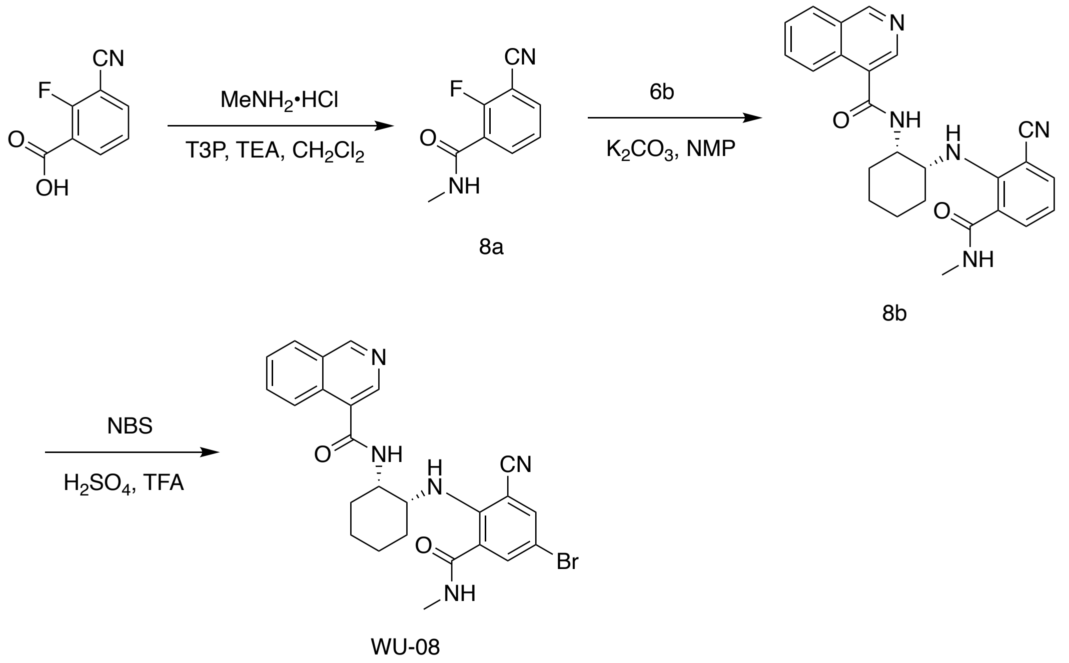


**Scheme S8. The synthesis route of compound WU-08.**

***Preparation of compound 3-cyano-2-fluoro-N-methylbenzamide (8a)***

To a solution of 3-cyano-2-fluoro-benzoic acid (10.0 g, 60.6 mmol) in $CH_2Cl_2$ (90 mL), T3P (57.8 g, 90.8 mmol, 54 mL, 50% purity), TEA (18.4 g, 182 mmol, 25.3 mL), and methanamine hydrochloride (4.91 g, 72.6 mmol) were added. The mixture was stirred at 25 °C for 1 hour. The reaction mixture was quenched by addition of $H_2O$ (30 mL) at 25 °C, and then extracted with $CH_2Cl_2$ (30 mL x 3). The organic layers were combined, washed with saturated aqueous NaCl (30 mL x 3), dried over $Na_2SO_4$, and filtered. The solvent was removed by evaporation under reduced pressure, and the residue was purified by flash silica gel chromatography to give compound **8a** as a yellow solid (6.60 g, 32.1% yield).

$^1$H NMR (400 MHz, $CD_3OD$): δ 7.95-8.02 (m, 1H), 7.86-7.91 (m, 1H), 7.43-7.47 (m, 1H), 2.94-2.95 (m, 3H). LC-MS: m/z = 179.2 $[M+H]^+$.

***Preparation of compound N-[(1S,2R)-2-[2-cyano-6-(methylcarbamoyl)anilino] cyclohexyl]isoquinoline-4-carboxamide (8b)***

To a solution of compound **8a** (3.37 g, 18.9 mmol) in NMP (50 mL), $K_2CO_3$ (5.23 g, 37.8 mmol) and N-[(1S,2R)-2-aminocyclohexyl]isoquinoline-4-carboxamide (compound **6b**, 5.10 g, 18.9 mmol) were added. After stirring at 120 °C for 12 hours, the reaction was quenched by adding $H_2O$ (30 mL) at 25 °C, and extracted with ethyl acetate (50 mL x 5). The organic layers were combined, washed with saturated aqueous NaCl (50 mL x 5), dried over $Na_2SO_4$, and filtered. The solvent was removed by evaporation under reduced pressure, and the residue was purified by flash silica gel chromatography to give compound **8b** as a yellow solid (5.40 g, 40.3% yield).

$^1$H NMR (400 MHz, $CD_3OD$): δ 9.27 (s, 1H), 8.47 (s, 1H), 8.09-8.15 (q, 2H), 7.77-7.81 (m, 1H), 7.69-7.74 (m, 1H), 7.63-7.67 (m, 1H), 7.55-7.60 (m, 1H), 6.70-6.74 (t, 1H), 4.76-4.79 (m, 1H), 4.57-4.59 (m, 1H), 2.81 (s, 3H), 1.83-1.93 (m, 4H), 1.65-1.78 (m, 2H), 1.54-1.62 (m, 2H). LC-MS: m/z = 428.3 $[M+H]^+$.

***Preparation of compound N-[(1S,2R)-2-[4-bromo-2-cyano-6-(methylcarbamoyl) anilino]cyclohexyl]isoquinoline-4-carboxamide (WU-08)***

To a solution of of compound **8b** (5.40 g, 12.6 mmol) in a mixture of $H_2SO_4$ (30 mL) and TFA (20 mL), NBS (2.25 g, 12.6 mmol) was added. After stirring at 20 °C for 1 hour, the reaction mixture was poured into water (50 mL) and the pH was adjusted to 7 using saturated aqueous $Na_2CO_3$. The mixture was then extracted with ethyl acetate (50 mL x 2). The organic layers were combined, washed with saturated aqueous NaCl (50 mL x 2), dried with anhydrous $Na_2SO_4$, and filtered. The solvent was removed by evaporation under reduced pressure, and the residue was purified by prep-HPLC to give compound **WU-08** as a yellow solid (2.80 g, 5.40 mmol, 42.7% yield, 97.66% purity).

[1]H NMR (400 MHz, $CD_3OD$): δ 9.28 (s, 1H), 8.54 (s, 1H), 8.13-8.15 (d, J = 8 Hz, 1H), 8.07-8.09 (d, J = 8 Hz, 1H), 7.78-7.84 (m, 1H), 7.69-7.77 (m, 3H), 4.83 (s, 1H), 4.58 (s, 1H), 2.78 (s, 3H), 1.81-1.93 (m, 4H), 1.67 (m, 2H), 1.53-1.63 (m, 2H). LC-MS: m/z = 506.3 $[M+H]^+$.

Synthesis of compounds WU-09 and WU-10

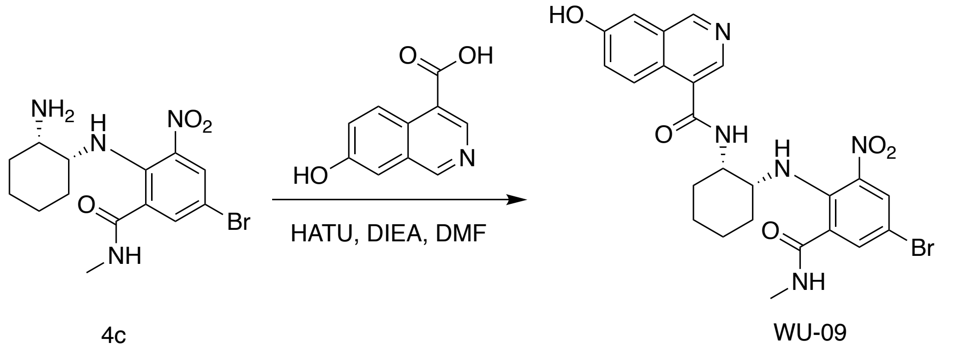


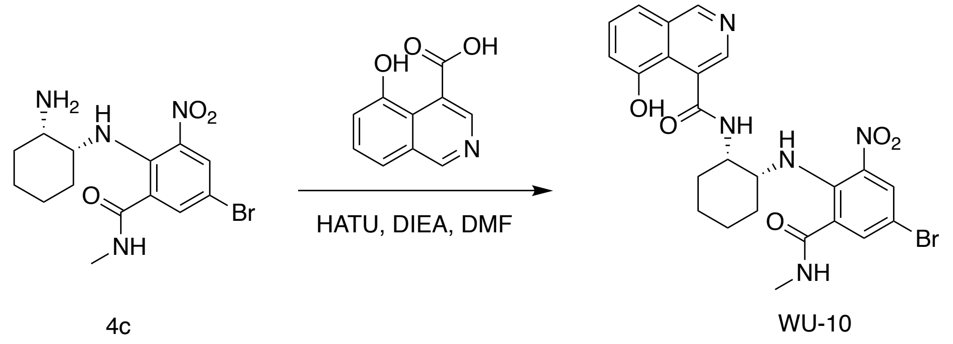


**Scheme S9. The synthesis routes of compounds WU-09 and WU-10.**

***Preparation of compound N-((1S,2R)-2-((4-bromo-2-(methylcarbamoyl)-6-nitrophenyl)amino)cyclohexyl)-7-hydroxyisoquinoline-4-carboxamide (WU-09)***

To a solution of 7-hydroxyisoquinoline-4-carboxylic acid (57 mg, 0.3 mmol) in anhydrous $CH_2Cl_2$ (5 mL), oxalyl chloride (42 mg, 0.33 mmol) was added, followed by the addition of 2-3 drops of anhydrous DMF. After stirring at room temperature for 1 hour, the solvent was removed by evaporation. The residue was dissolved in anhydrous $CH_2Cl_2$ (5 mL) and added to a solution of compound 2-(((1R, 2S)-2-aminocyclohexyl) amino)-5-bromo-N-methyl-3-nitrobenzamide (compound **4c**, 0.3 mmol) and DIEA (116 mg, 0.9 mmol) in $CH_2Cl_2$. The reaction mixture was stirred at room temperature overnight, and then extracted with $CH_2Cl_2$. The organic layers were combined, washed with saturated aqueous NaCl, dried over anhydrous $Na_2SO_4$. The solvent was removed, and the residue was purified by flash silica gel chromatography to give compound **WU-11** as a yellow solid (12 mg, 7% yield).

$^1$H NMR (500 MHz, $CDCl_3$): δ 10.62 (s, 1H), 9.10 (s, 1H), 8.53 (s, 1H), 8.02 (d, $J$ = 2.4 Hz, 1H), 7.87 (d, $J$ = 10.9 Hz, 1H), 7.56 (m, 2H), 7.49 (m, 1H), 7.33-7.27 (m, 2H), 6.90 (d, $J$ = 4.5 Hz, 1H), 4.33 (d, $J$ = 7.2 Hz, 1H), 3.99 (d, $J$ = 5.8 Hz, 1H), 2.98 (d, $J$ = 4.8 Hz, 3H), 2.03-1.48 (m, 8H). ESI-MS: $m/z$ = 542.00 $[M+H]^+$.

***Preparation of compound N-((1S,2R)-2-((4-bromo-2-(methylcarbamoyl)-6-nitrophenyl)amino)cyclohexyl)-5-hydroxyisoquinoline-4-carboxamide (WU-10)***

To a solution of 5-hydroxyisoquinoline-4-carboxylic acid (28 mg, 0.15 mmol) in anhydrous $CH_2Cl_2$ (5 mL), oxalyl chloride (21 mg, 0.165 mmol) was added, followed by the addition of 2-3 drops of anhydrous DMF. After stirring at room temperature for 2 hours, the solvent was removed by evaporation under reduced pressure. The residue was dissolved in anhydrous $CH_2Cl_2$ (5 mL) and added to a solution of 2-(((1R, 2S)-2-aminocyclohexyl) amino)-5-bromo-N-methyl-3-nitrobenzamide (compound **4c**, 0.14 mmol) and DIPEA (54 mg, 0.42 mmol) in $CH_2Cl_2$. The reaction mixture was stirred at room temperature overnight, and then extracted with $CH_2Cl_2$. The organic layers were combined, washed with saturated aqueous NaCl, dried over anhydrous $Na_2SO_4$. The solvent was removed, and the residue was purified by flash silica gel chromatography to give compound **WU-10** as a yellow solid (17 mg, 22% yield).

$^{1}H$ NMR (500 MHz, $CDCl_3$): δ 10.66 (s, 1H), 9.13 (s, 1H), 8.56 (s, 1H), 8.02 (s, 1H), 7.88 (d, $J$ = 10.2 Hz, 1H), 7.56 (m, 3H), 7.35 (m, 2H), 6.91 (s, 1H), 4.32 (s, 1H), 3.98 (s, 1H), 2.98 (d, $J$ = 4.3 Hz, 3H), 2.03-1.46 (m, 8H). ESI-MS: $m/z$ = 542.00 $[M+H]^+$.

Synthesis of compounds WU-11 and WU-12

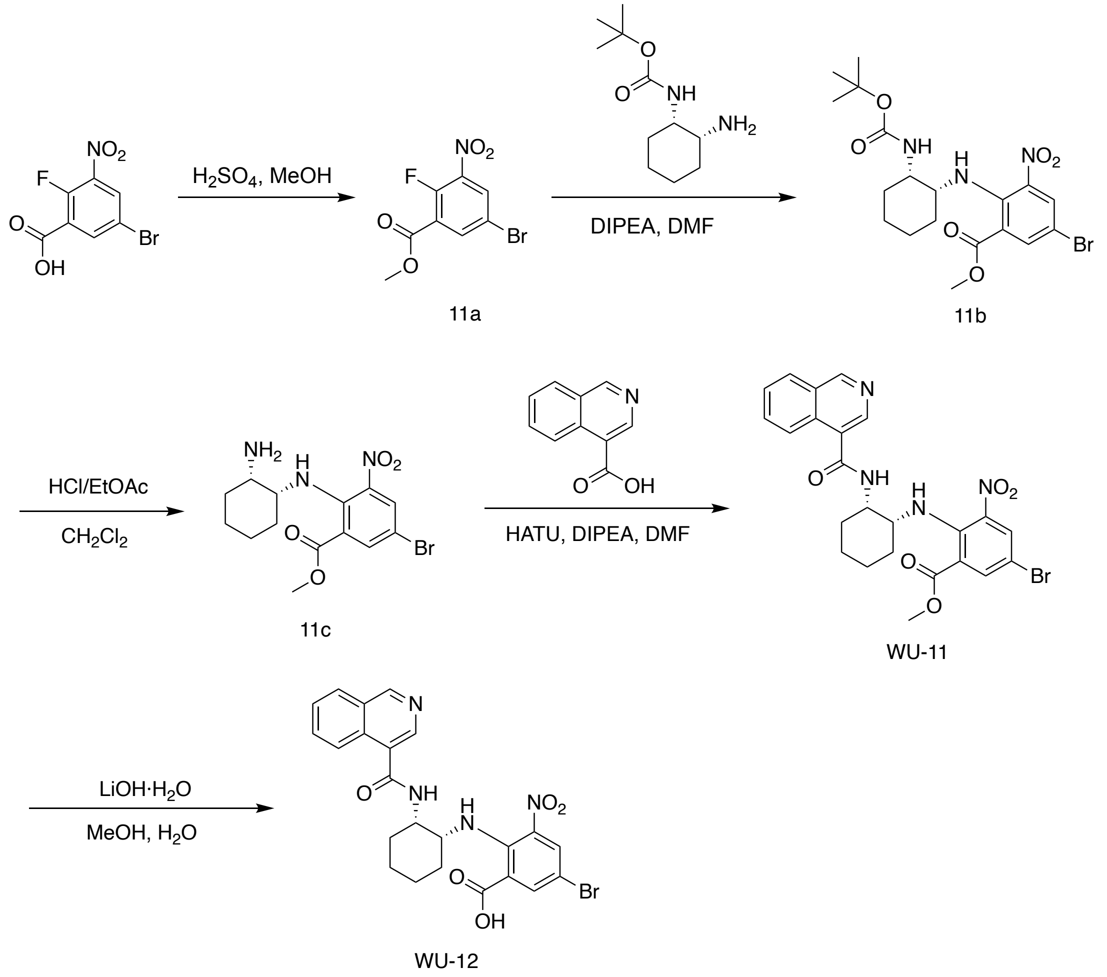


**Scheme S10. The synthesis routes of compounds WU-11 and WU-12.**

***Preparation of compound methyl-5-bromo-2-fluoro-3-nitrobenzoate (11a)***

A solution of 5-bromo-2-fluoro-3-nitrobenzoic acid (1 g, 3.80 mmol) in MeOH (20 mL) was stirred at room temperature. After the addition of conc. $H_2SO_4$ (0.5 mL), the reaction was heated to reflux for 3 hours. Then organic solvent was removed under reduced pressure and brine was added and the mixture was extracted with ethyl acetate. The organic layer was dried over anhydrous $Na_2SO_4$ and concentrated under vacuum. The residue was purified by chromatography using petroleum ether/ethyl acetate (v/v = 4/1). Compound **11a** was obtained as a yellow solid (0.71 g, 67.2% yield).

$^1$H NMR (500 MHz, $CDCl_3$): δ 8.31 (m, 2H), 3.99 (s, 3H, $CH_3$).

***Preparation of compound methyl 5-bromo-2-(((1R,2S)-2-((tert-butoxycarbonyl) amino)cyclohexyl)amino)-3-nitrobenzoate (11b)***

To a solution of compound **11a** (0.5 g, 1.81 mmol) in DMF (10 mL), tert-butyl ((1S,2R)-2-aminocyclohexyl)carbamate (0.47 g, 2.17 mmol) and DIPEA (0.9 mL, 5.43 mmol) were added with stirring at room temperature. The reaction was warmed to 80 °C and stirred for 16 hours. Then the mixture was extracted with ethyl acetate three times and the organic layers were combined and washed with saturated aqueous NaCl. The organic layer was dried over anhydrous $Na_2SO_4$ and concentrated under vacuum. Without further purification, compound **11b** was obtained as a yellow solid.

$^1$H NMR (500 MHz, $CDCl_3$): δ 8.36 (s, 1H), 8.11 (s, 1H), 8.04 (s, 1H), 4.58 (d, J = 7.2 Hz, 1H), 3.92 (s, 3H), 3.76 (d, J = 10.6 Hz, 1H), 3.50 (s, 1H), 1.60 (m, 6H), 1.51-1.40 (m, 4H), 1.37 (s, 9H).

***Preparation of compound methyl 2-(((1R,2S)-2-aminocyclohexyl)amino)-5-bromo-3-nitrobenzoate (11c)***

To a solution of compound **11b** (201 mg, 0.43 mmol) in anhydrous $CH_2Cl_2$ (6 mL) with stirring at room temperature, HCl (6 mL, 3M in ethyl acetate) was added. The reaction was stirred at room temperature for 1 hour. The mixture was concentrated under vacuum to remove the solvent. The residue was used in next step reaction without further purification.

***Preparation of compound methyl 5-bromo-2-(((1R,2S)-2-(isoquinoline-4-carboxamido)cyclohexyl)amino)-3-nitrobenzoate (WU-11)***

To a solution of isoquinoline-4-carboxylic acid (compound **6a**, 74 mg, 0.43 mmol) in 5 mL of DMF, HATU (245 mg, 0.64 mmol) and DIPEA (167 mg, 1.29 mmol) were added with stirring at room temperature. The reaction was monitored by thin-layer chromatography (TLC). After the reaction was completed, water was added, and compound **WU-11** was precipitated as a yellow solid. The yellow solid was filtered and dried to give 0.28 g **WU-11**.

$^1$H NMR (500 MHz, $CDCl_3$): δ 9.25 (s, 1H), 8.63 (s, 1H), 8.47 (d, J = 10.5 Hz, 1H), 8.30 (d, J = 8.5 Hz, 1H), 8.10 (d, J = 2.4 Hz, 1H), 7.98 (m, 2H), 7.78 (t, J = 7.7 Hz, 1H), 7.66 (t, J = 7.5 Hz, 1H), 6.42 (d, J = 8.1 Hz, 1H), 4.37 (m, 1H), 3.82 (s, 3H), 3.71 (m, 1H), 1.81-1.46 (m, 8H). ESI-MS: m/z = 527.29 $[M+H]^+$.

***Preparation of compound 5-bromo-2-(((1R,2S)-2-(isoquinoline-4-carboxamido)cyclohexyl)amino)-3-nitrobenzoic acid (WU-12)***

$LiOH{\cdot}H_2O$ (36 mg, 0.87 mmol) was added to a solution of compound **WU-11** (150 mg, 0.29 mmol) in a mixture of 10 mL of MeOH (10 mL) and 1 mL of water. The reaction was stirred at room temperature for 2 hours. Then the organic solvent was removed under vacuum and the residue was dissolved in 2 mL of water. After the pH was adjusted to 5 using 2N HCl, the precipitate was collected by filtration and dried to give **WU-12** as a yellow solid (100 mg). $^1$H NMR (500 MHz, $CDCl_3$): δ 9.04 (s, 1H), 8.71 (s, 1H), 8.34 (d, J = 8.0 Hz, 1H), 8.08 (s, 1H), 7.87 (m, 3H), 7.66 (t, J = 7.5 Hz, 1H), 7.43 (s, 1H), 4.41 (s, 1H), 3.70 (s, 1H), 1.87-1.52 (m, 8H). ESI-MS: m/z = 518.28 $[M+H]^+$.

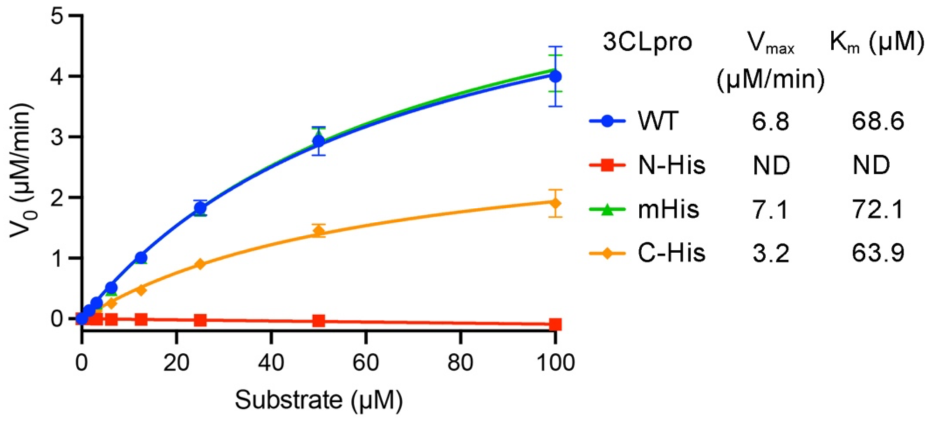


**Fig. S1 | Enzyme activity of wild type 3CLpro and 6x His-tagged 3CLpro of SARS-CoV-2.**

WT is the 3CLpro of SARS-CoV-2 with no additional residues at its N or C terminus. mHis is the 3CLpro of SARS-CoV-2 with a 6x His tag inserted between residues Arg222 and Phe223. N-His and C-His are the 3CLpro of SARS-CoV-2 with a 6x His tag at its N-terminus and C-terminus, respectively. The enzyme activity was evaluated using a fluorescence resonance energy transfer (FRET)-based assay (see "Materials and Methods"). The data represent the mean ± SD of four independent measurements.

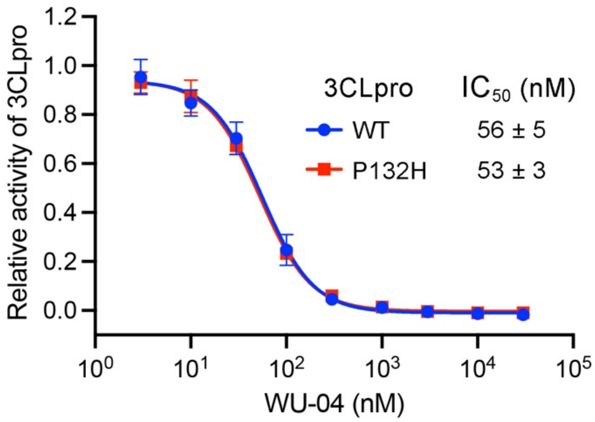


**Fig. S2 | Inhibition of the 3CLpro of the Omicron variant of SARS-CoV-2.**

The ability of compound WU-04 to inhibit the enzyme activity of the wild-type (WT) 3CLpro and that of the 3CLpro of the SARS-CoV-2 Omicron variant which carries a P132H mutation was evaluated using the fluorescent substrate Dabcyl-KTSAVLQSGFRKME-Edans. The data represent the mean ± SD of three independent measurements.

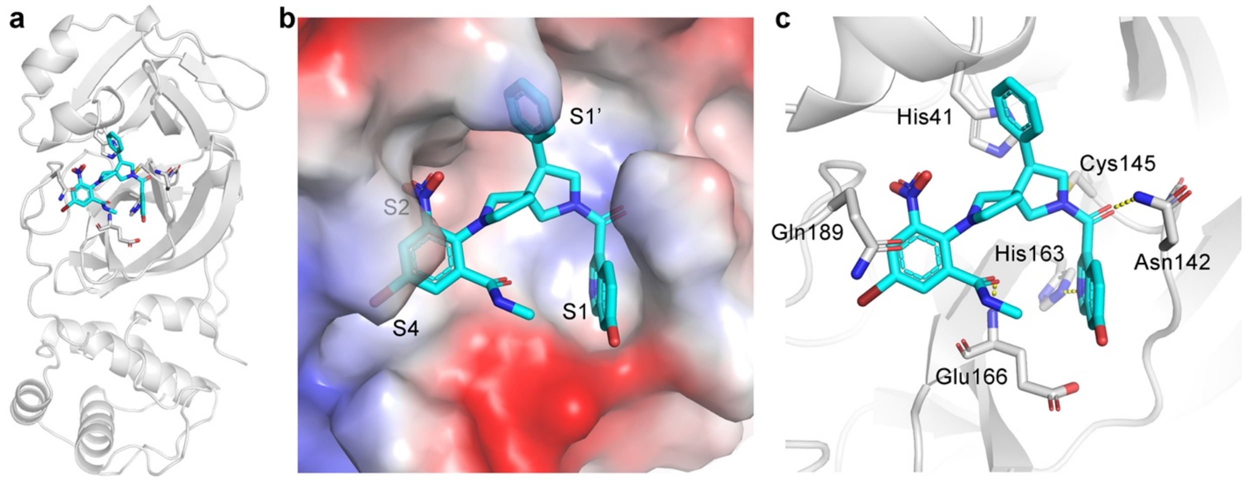


**Fig. S3 | The crystal structure of SARS-CoV-2 3CLpro in complex with WU-02. a**, The overall structure of the 3CLpro/WU-02 complex. The inhibitor WU-02 is colored cyan. **b**, The binding mode of WU-02 in the catalytic pocket of 3CLpro. The isoquinoline ring of WU-02 binds to site S1. The 6-nitro and 4-bromo groups of the bromophenyl ring of WU-02 are docked into sites S2 and S4, respectively. The phenyl group of WU-02 extents to site S1'. **c,** The interactions between WU-02 and the residues in 3CLpro. The yellow dash lines represent hydrogen bonds.

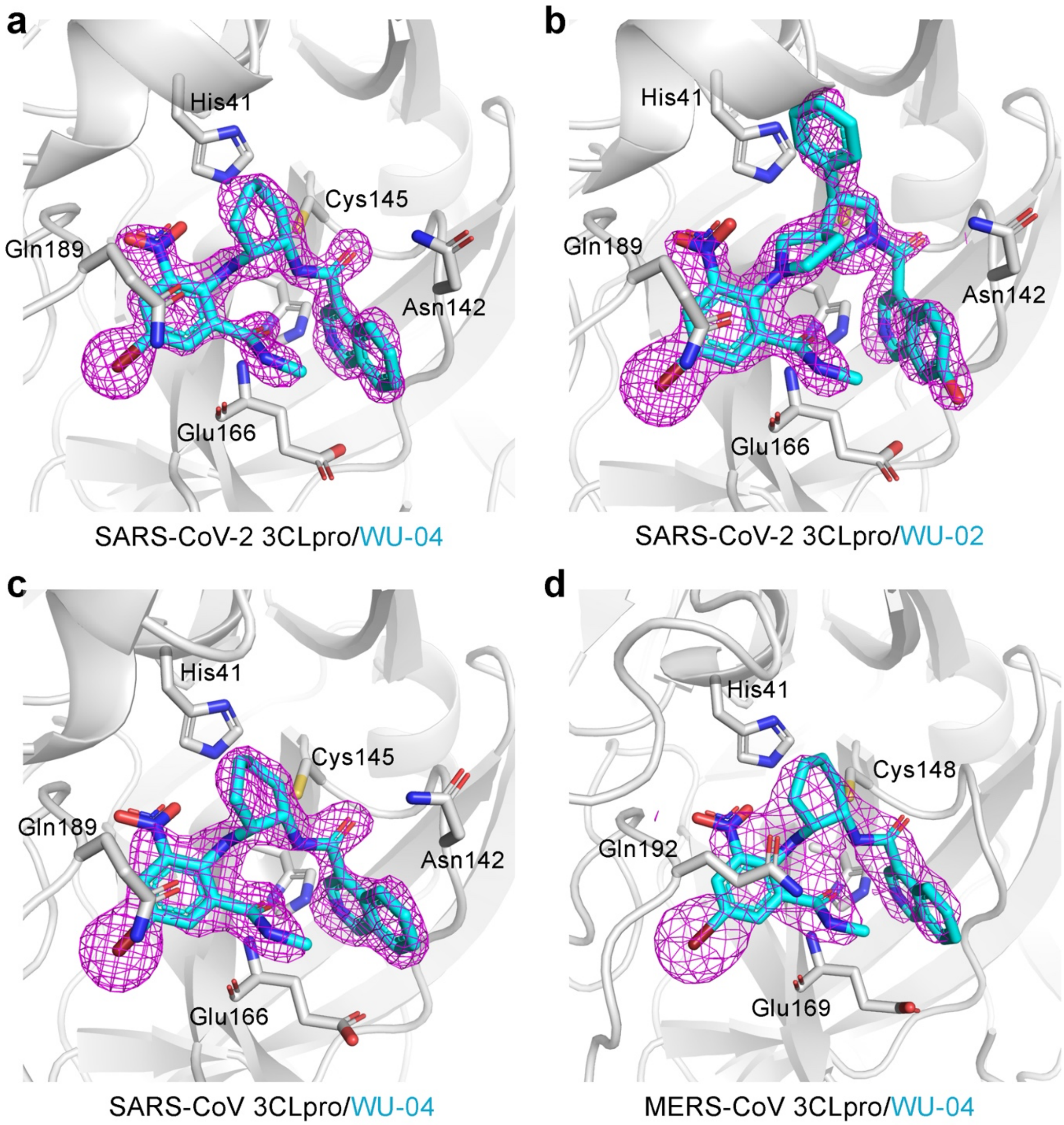


**Fig. S4 | The composite omit maps the non-covalent inhibitor WU-02 and WU-04 in the 3CLpro crystal structures. a**, **c** and **d**, The composite omit maps of WU-04 in the crystal structures of the SARS-CoV-2 3CLpro/WU-04 complex (**a**), the SARS-CoV 3CLpro/WU-04 complex (**c**), and the MERS-CoV 3CLpro/WU-04 complex (**d**). **b**, The composite map of WU-02 in the crystal structure of the SARS-CoV-2 3CLpro/WU-02 complex. All the maps were generated using the composite_omit_map program **(20)** in PHENIX. The 2mFo-DFc electron density maps were contoured at 2.0 σ and colored magenta.

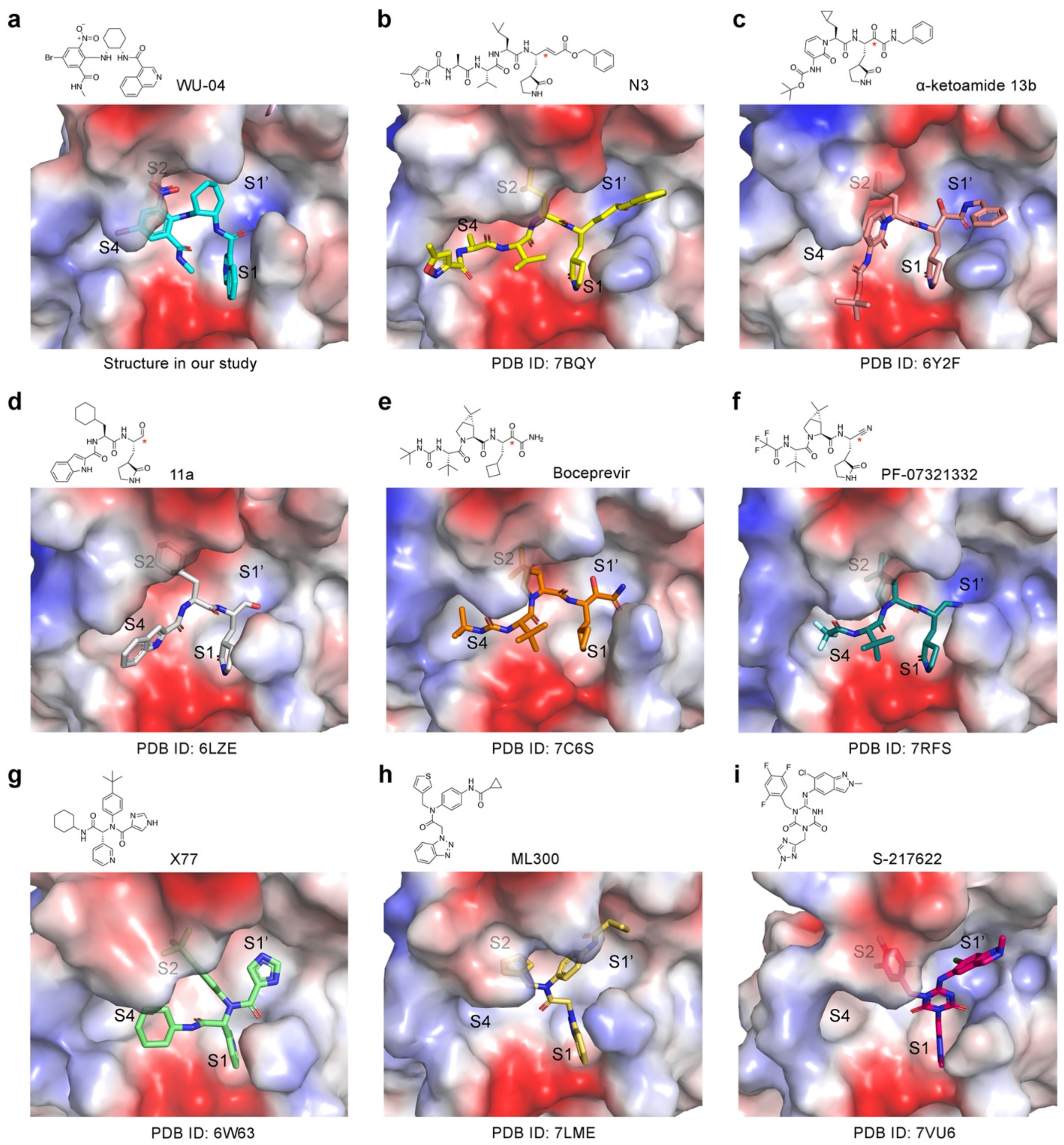


**Fig. S5 | Comparison of the binding mode of WU-04 to 3CLpro with that of four covalent inhibitors and a non-covalent inhibitor of SARS-CoV-2 3CLpro. a**, The binding mode of WU-04 to SARS-CoV-2 3CLpro. **b-f**, The binding modes of five covalent inhibitors to SARS-CoV-2 3CLpro. The reaction site in each inhibitor is indicated by a red asterisk. Compound N3 (**b**) was first developed as a pan-inhibitor of coronavirus 3CLpro (21), and then found to be also an SRAS-

CoV-2 3CLpro inhibitor (2). Compound α-ketoamide 13b (**c**) was designed on the basis of a covalent inhibitor of MERS-CoV and SARS-CoV (22). Compound 11a (**d**) was designed by analyzing reported SARS-CoV 3CLpro inhibitors (23). Boceprevir (**e**) is an inhibitor of the HCV NS3 protease and was identified to be also an inhibitor of SARS-CoV-2 3CLpro (24). PF-07321332 (**f**) was developed by Pfizer as an oral SARS-CoV-2 3CLpro inhibitor (25). **g**-**i**, Binding mode of three non-covalent inhibitors to SARS-CoV-2 3CLpro. The crystal structure of compound X77 (**g**) in complex with SARS-CoV-2 3CLpro has been deposited in Protein Data Bank, but the research has not been published (26). The structure of compound X77 is similar to compound ML188 that was previously developed as a non-covalent inhibitor of SARS-CoV 3CLpro (27). Compound ML300 (**h**) was first identified as a non-covalent inhibitor of SARS-CoV 3CLpro (28), then it was also showed weak activity to inhibit SARS-CoV-2 3CLpro (29); a series of ML300 derivatives was synthesized, and the most potent one, compound CCF0058981, showed an $EC_{50}$ of about 0.5 μM in an anti-SARS-CoV-2 assay in Vero E6 ACE2 cells (29). Compound S-217622 was a non-covalent oral SARS-CoV-2 3CLpro inhibitor developed by Shionogi & Co., Ltd. (30).

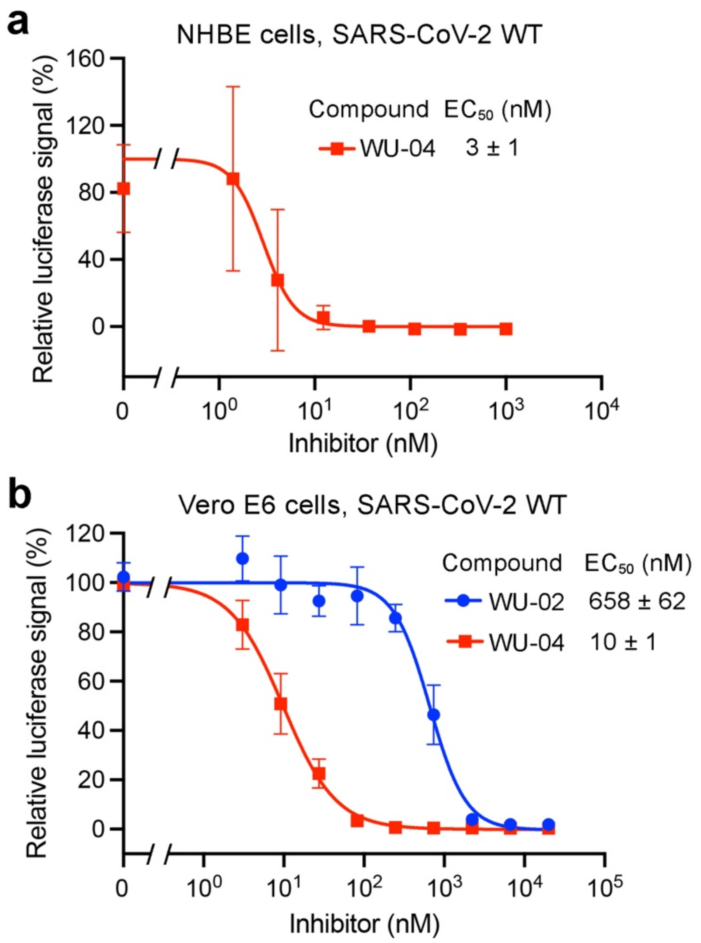


**Fig. S6 | The non-covalent 3CLpro inhibitors effectively inhibited the replication of SARS-CoV-2 in NHBE cells (a) and in Vero E6 cells (b).** The data represent the mean ± SD of six (**a**) or four (**b**) independent measurements.

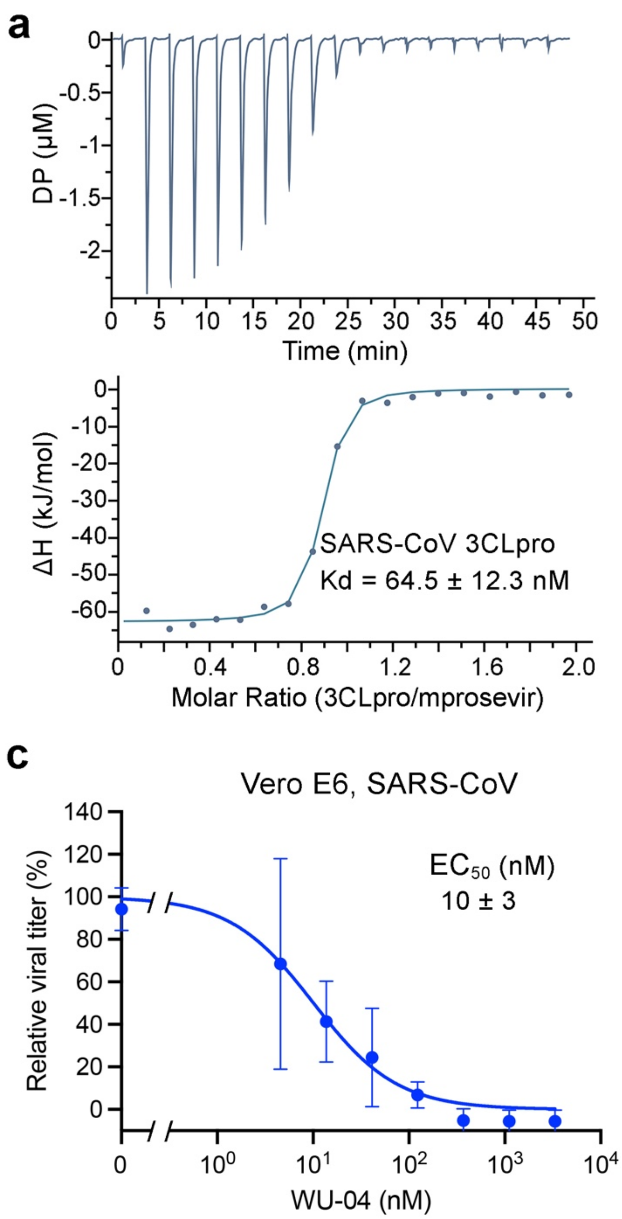


**Fig. S7 | WU-04 bound to and inhibited the 3CLpro of SARS-CoV and MERS-CoV. a** and **b**, The binding affinity ($K_d$) between WU-04 and SARS-CoV 3CLpro (**a**) and that between WU-04 and MERS-CoV 3CLpro (**b**) were measured using isothermal titration calorimetry (ITC). **c** and **d**, The antiviral activity of WU-04 against SARS-CoV (**c**) or against MERS-CoV (**d**) was measured in Vero E6 cells. The data represents the mean ± SD of three independent measurements.

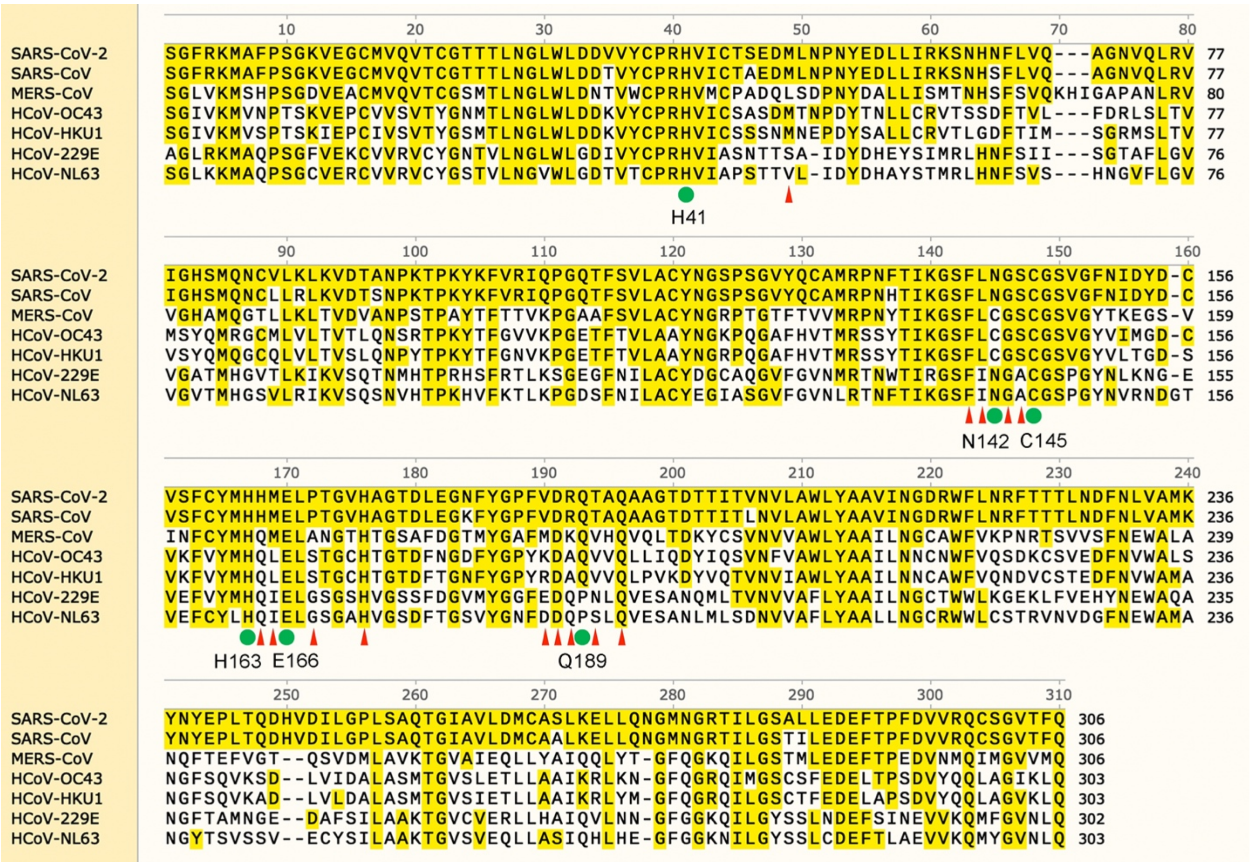

**Fig. S8 | Alignment of the human coronavirus 3CLpro protein sequences.**

The protein sequences of 3CLpro from SARS-CoV-2 (NCBI accession number: YP_009725301.1), SARS-CoV (NCBI accession number: NP_828863.1), MERS-CoV (NCBI accession number: YP_009047217.1), HCoV-OC43 (NCBI accession number: YP_009555250.1), HCoV-HKU1 (NCBI accession number: YP_459936.1), HCoV-229E (NCBI accession number: NP_073549.1, residues 2966 to 3267), and HCoV-NL63 (NCBI accession number: YP_003766.2, residues 2940 to 3242) were aligned using SnapGene (version 5.2.4). The blue dots indicate the catalytic residues and residues in 3CLpro that are important for binding with WU-04. The red triangles indicate other residues in 3CLpro that are within 5 Å of WU-04 according to the crystal structure of the SARS-CoV-2 3CLpro/WU-04 complex.

**a**

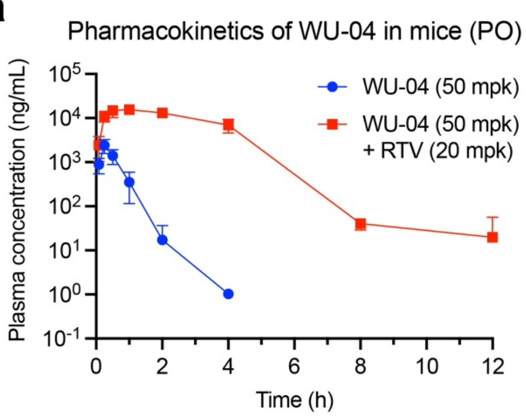


**b**

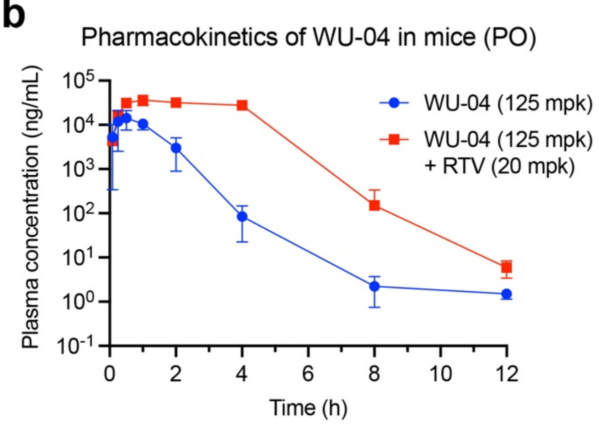


**c**

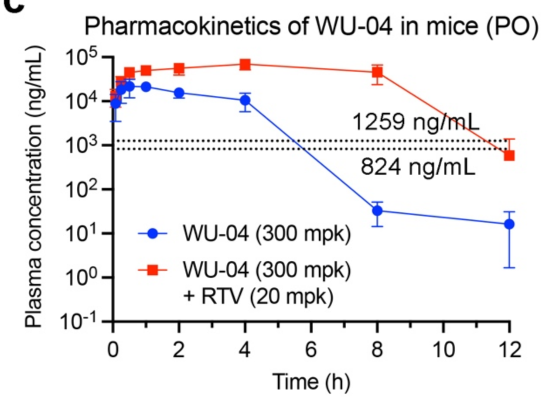


**d**

| PK Parameters in Mice | $C_{max}$ (ng/mL) | $T_{1/2}$ (h) | $AUC_{0-inf}$ (ng.h/mL) |
|---|---|---|---|
| **WU-04 (50 mpk)** | 2417 ± 835 | 0.25 ± 0.01 | 1260 ± 414 |
| **WU-04 (50 mpk) + RTV (20 mpk)** | 17423 ± 2858 | 1.54 ± 0.50 | 51051 ± 7205 |
| **WU-04 (125 mpk)** | 17185 ± 6465 | 1.49 ± 1.24 | 18461 ± 7571 |
| **WU-04 (125 mpk) + RTV (20 mpk)** | 37936 ± 8107 | 2.30 ± 0.45 | 139539 ± 27677 |
| **WU-04 (300 mpk)** | 25381 ± 8630 | 2.02 ± 1.59 | 69796 ± 12780 |
| **WU-04 (300 mpk) + RTV (20 mpk)** | 69934 ± 18571 | 4.00 ± 2.12 | 488615 ± 155600 |

**e**

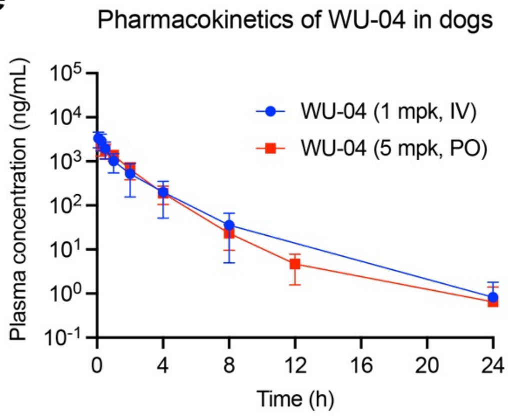


**f**

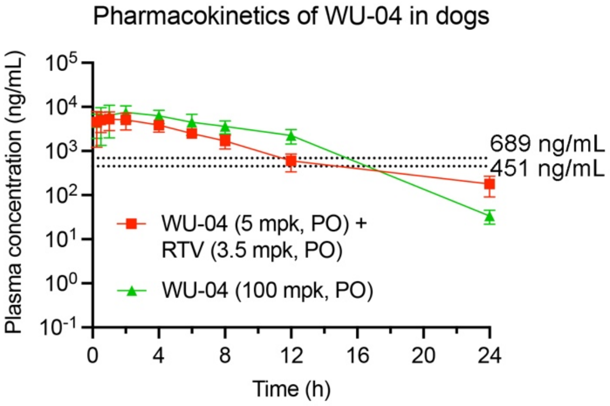


**g**

| PK Parameters in Dogs | WU-04 (1 mpk, IV) | WU-04 (5 mpk, PO) | WU-04 (5 mpk, PO) + RTV (3.5 mpk, PO) | WU-04 (100 mpk, PO) |
|---|---|---|---|---|
| **$C_{max}$ (ng/mL)** | 3542 ± 1384 | 2200 ± 889 | 5320 ± 2335 | 8163 ± 3105 |
| **$T_{1/2}$ (h)** | 1.90 ± 0.80 | 1.68 ± 0.52 | 4.83 ± 0.85 | 2.28 ± 0.29 |
| **$Vd_{ss}$ (L/kg)** | 0.45 ± 0.11 | / | / | / |
| **Cl(mL/min/kg)** | 5.55 ± 3.79 | / | / | / |
| **$AUC_{0-inf}$ (ng.h/mL)** | 4052 ± 2190 | 3670 ± 1332 | 38494 ± 13294 | 62010 ± 19721 |
| **Bioavailability (%)** | / | 18.1 | / | / |

**Fig. S9 | Pharmacokinetics of WU-04 in male BALB/c mice and in male Beagle dogs.**

**a** to **d**, After oral administration (PO) of WU-04 alone or together with ritonavir (RTV, 20 mpk), the blood samples of the mice were collected and processed for plasma by centrifugation. The plasma concentrations of WU-04 were determined through LC-MS/MS analysis. The data represents the mean ± SD (n = 5). **e** to **g**, WU-04 was administered alone or together with RTV (3.5 mpk) by intravenous injection (IV) or PO route, then the blood samples of the dogs were collected and the plasma concentrations of WU-04 were determined through LC-MS/MS analysis. The data represents the mean ± SD (in fig. S9**e**: n = 4; in fig S9**f**: n = 3). The two dash lines in **c** (and in **d**) indicate the total $EC_{90}$ values calculated based on the plasma unbound ratio of WU-04 (CD-1 mouse plasma: 2.3%; Beagle dog plasma: 4.2%) and the $EC_{90}$ values of WU-04 in A549-hACE2 cells (Fig. 3a) and in Calu-3 cells (Fig. 3c), respectively.

### Table S1 | Data collection and refinement statistics.

| | SARS-CoV-2 3CLpro/WU-02 | SARS-CoV-2 3CLpro/WU-04 | SARS-CoV 3CLpro/WU-04 | MERS-CoV 3CLpro/WU-04 |
|---|---|---|---|---|
| **Wavelength** | 1.541 | 1.541 | 1.541 | 1.541 |
| **Resolution range** | 29.51 - 1.9 (1.968 - 1.9)[a] | 29 - 1.83 (1.895 - 1.83)[a] | 30.76 - 1.99 (2.061 - 1.99)[a] | 29.18 - 2.98 (3.086 - 2.98)[a] |
| **Space group** | I 1 2 1 | P 21 21 21 | I 1 2 1 | P 1 21 1 |
| **Unit cell:** ***a*, *b*, *c* (Å)** | 54.3613, 80.3185, 87.2808 | 54.8299, 67.8594, 167.557 | 53.4961 82.6766 107.454 | 62.7555 112.952 92.3545 |
| **Unit cell:** ***α*, *β*, *γ* (°)** | 90, 96.9635, 90 | 90, 90, 90 | 90, 103.64, 90 | 90, 91.3159, 90 |
| **Total reflections** | 141388 (9747) | 220465 (14626) | 342964 (26236) | 124328 (13212) |
| **Unique reflections** | 29294 (2916) | 55659 (5444) | 31238 (3133) | 25971 (2568) |
| **Multiplicity** | 4.8 (3.3) | 4.0 (2.7) | 11.0 (8.4) | 4.8 (5.1) |
| **Completeness (%)** | 99.61 (99.08) | 99.27 (98.16) | 99.46 (99.62) | 98.42 (99.38) |
| **Mean I/sigma(I)** | 18.24 (3.17) | 21.70 (1.96) | 33.16 (5.12) | 10.58 (3.72) |
| **Wilson B-factor** | 21.01 | 13.76 | 21.89 | 36.42 |
| **R-merge** | 0.1339 (0.3152) | 0.125 (0.3773) | 0.1236 (0.4076) | 0.2467 (0.4305) |
| **R-meas** | 0.1488 (0.3763) | 0.1428 (0.4617) | 0.1295 (0.4344) | 0.2769 (0.4803) |
| **R-pim** | 0.06396 (0.2016) | 0.0677 (0.26) | 0.03849 (0.149) | 0.1235 (0.2094) |
| **CC1/2** | 0.986 (0.861) | 0.984 (0.861) | 0.997 (0.864) | 0.947 (0.866) |
| **CC*** | 0.997 (0.962) | 0.996 (0.962) | 0.999 (0.963) | 0.986 (0.963) |
| **Reflections used in refinement** | 29293 (2916) | 55656 (5443) | 31084 (3121) | 25960 (2567) |
| **Reflections used for R-free** | 1453 (145) | 1907 (187) | 1477 (149) | 1181 (118) |
| **R-work** | 0.1959 (0.2219) | 0.2244 (0.3710) | 0.1982 (0.2265) | 0.2342 (0.2930) |
| **R-free** | 0.2322 (0.2608) | 0.2731 (0.4192) | 0.2379 (0.2530) | 0.2711 (0.3346) |
| **CC (work)** | 0.952 (0.893) | 0.943 (0.804) | 0.963 (0.884) | 0.925 (0.801) |
| **CC (free)** | 0.942 (0.856) | 0.939 (0.658) | 0.951 (0.843) | 0.886 (0.772) |
| **Number of non-hydrogen atoms** | 2664 | 5313 | 2666 | 9384 |
| **macromolecules** | 2368 | 4697 | 2371 | 9248 |
| **ligands** | 42 | 74 | 34 | 136 |
| **solvent** | 254 | 542 | 261 | |
| **Protein residues** | 306 | 607 | 306 | 1216 |

| | | | | |
|---|---|---|---|---|
| **RMS (bonds)** | 0.007 | 0.007 | 0.003 | 0.006 |
| **RMS (angles)** | 0.86 | 0.94 | 0.64 | 1.02 |
| **Ramachandran favored (%)** | 98.68 | 98.01 | 98.03 | 97.93 |
| **Ramachandran allowed (%)** | 1.32 | 1.66 | 1.97 | 2.07 |
| **Ramachandran outliers (%)** | 0.00 | 0.33 | 0.00 | 0.00 |
| **Rotamer outliers (%)** | 1.14 | 0.38 | 0.76 | 0.20 |
| **Clashscore** | 2.96 | 3.63 | 2.96 | 6.89 |
| **Average B-factor** | 26.04 | 15.61 | 25.99 | 36.32 |
| **macromolecules** | 25.66 | 15.15 | 25.50 | 36.25 |
| **ligands** | 25.19 | 11.74 | 23.21 | 41.19 |
| **solvent** | 29.69 | 20.15 | 30.85 | |

[a] Statistics for the highest-resolution shell are shown in parentheses.

**Table S2. Inhibition of metabolism of WU-04 in human liver microsomes by CYP isoform-selective chemical inhibitors***

| %Inhibition of metabolism of WU-04 (Depletion Method) | | | | | | |
|---|---|---|---|---|---|---|
| **CYP1A2** | **CYP2B6** | **CYP2C8** | **CYP2C9** | **CYP2C19** | **CYP2D6** | **CYP3A** |
| 0.0 | -2.4 | 9.8 | -0.1 | -6.7 | -3.1 | 84.4 |

*The inhibitors used for CYP1A2, CYP2B6, CYP2C8, CYP2C9, CYP2C19, CYP2D6, and CYP3A were α-Naphthoflavone, Ticlopidine, Montelukast, Sulfaphenazole, (+)-N-3-benzylnirvanol, Quinidine, and Ketoconazole, respectively.

### Table S3. Inhibition of metabolism of WU-04 in liver microsomes by ritonavir (RTV)*

| Sample Name | Human Liver Microsomes (HLM) | | | | | | CD-1 Mouse Liver Microsomes (MLM) | | | | | | Beagle Dog Liver Microsomes (DLM) | | | | | |
|---|---|---|---|---|---|---|---|---|---|---|---|---|---|---|---|---|---|---|
| | $R^2$ | $T_{1/2}$ (min) | $CL_{int(mic)}$ (μL/min/mg) | $CL_{int(liver)}$ (mL/min/kg) | Remaining (T=60 min) | Remaining (NCF=60min) | $R^2$ | $T_{1/2}$ (min) | $CL_{int(mic)}$ (μL/min/mg) | $CL_{int(liver)}$ (mL/min/kg) | Remaining (T=60 min) | Remaining (NCF=60min) | $R^2$ | $T_{1/2}$ (min) | $CL_{int(mic)}$ (μL/min/mg) | $CL_{int(liver)}$ (mL/min/kg) | Remaining (T=60 min) | Remaining (NCF=60min) |
| WU-04 | 1.0000 | 0.9 | 1531.1 | 1378.0 | 0.0% | 97.8% | 1.0000 | 0.9 | 1557.3 | 6166.8 | 0.1% | 98.5% | 0.9320 | 2.3 | 595.8 | 857.9 | 0.3% | 97.3% |
| WU-04 + RTV | 0.9212 | >145 | <9.6 | <8.6 | 74.1% | 79.8% | 0.9418 | 87.1 | 15.9 | 63.0 | 60.5% | 104.6% | 0.9890 | 69.7 | 19.9 | 28.6 | 54.6% | 82.6% |

* **$R^2$**: correlation coefficient of the linear regression for the determination of kinetic constant; **$T_{1/2}$**: half life; **$CL_{int(mic)}$**: intrinsic clearance, $CL_{int(mic)}$ = 0.693/$T_{1/2}$/mg microsome protein per mL; **$CL_{int(liver)}$** = $CL_{int(mic)}$ * mg microsomal protein/g liver weight * g liver weight/kg body weight; **NCF** (abbreviation of no co-factor): no NADPH is added to NCF samples (replaced by buffer) during the 60 minute incubation.
